# The single-PDZ domain protein TXBP-3 regulates contractility in the *C. elegans* spermatheca

**DOI:** 10.64898/2026.10.05.756805

**Authors:** Maria Khalid, Erin J. Cram

## Abstract

In the *C. elegans* reproductive system, oocyte transit through the spermatheca relies on parallel calcium and Rho-dependent signaling pathways to regulate actomyosin contractility. Here, we identify the conserved single-PDZ domain protein TXBP-3/TAX1BP3 as a dosage-dependent regulator of spermathecal contractility. TXBP-3 is expressed in contractile tissues, including the spermatheca and spermathecal-uterine (sp-ut) valve. Animals lacking *txbp-3* do not display overt phenotypes or defects in ovulation. However, in nematodes expressing the genetically encoded Ca^2+^ sensor GCaMP, deletion of *txbp-3* results in more rapid transit of embryos through the spermatheca. Conversely, tissue-specific overexpression of TXBP-3 results in slower transit times due to hypercontraction of the spermathecal neck and sp-ut valve, causing oocyte severing and failures of embryos to exit the spermatheca. We find TXBP-3 promotes the active phosphorylated state of the Ca^2+^/calmodulin-dependent protein kinase II (CaMKII) UNC-43. When TXBP-3 is overexpressed or *unc-43* is depleted, ovulations in which the embryo fails to exit show disrupted Ca^2+^ signaling. Additionally, TXBP-3 functions independently of the RhoGEF RHGF-1 and in parallel with the RhoGAP SPV-1 to maintain the alignment of actin fiber bundles and protect embryo structural integrity. Together, our results establish TXBP-3 as a regulator of contractility, kinase activation, and cytoskeletal architecture in contractile tissues.

## Introduction

Regulation of cellular contraction and relaxation is essential for the function of epithelial and endothelial tubes, which transport fluids and gases throughout the body. Disruptions in these processes are linked to diseases such as hypertension and asthma (Fabry & Fredberg, 2007; Orr et al., 2006). The *Caenorhabditis elegans* spermatheca, the site of oocyte fertilization, is a contractile tissue and serves as a model for studying the coordination of contractility (Kariya et al., 2004; Wirshing & Cram, 2018). This organ consists of 24 smooth muscle-like cells that form an entry neck and a central bag, attached to the syncytial spermathecal-uterine (sp-ut) valve (Hirsh et al., 1976; Hubbard & Greenstein, 2000) (Fig. 1A). During ovulation, the entering oocyte stretches the tissue, triggering a Ca^2+^ signaling wave and actomyosin-mediated contraction, which pushes the fertilized embryo into the uterus (McCarter et al., 1997; Yamamoto et al., 2006; Kovacevic et al., 2013; Sethi et al., 2017). Poor coordination of these contractions and relaxations can lead to fragmented, trapped embryos or ruptured oocytes, which can cause reproductive failure (Kovacevic & Cram, 2010; Tan & Zaidel-Bar, 2015; Bouffard et al., 2019).

**Figure 1.**
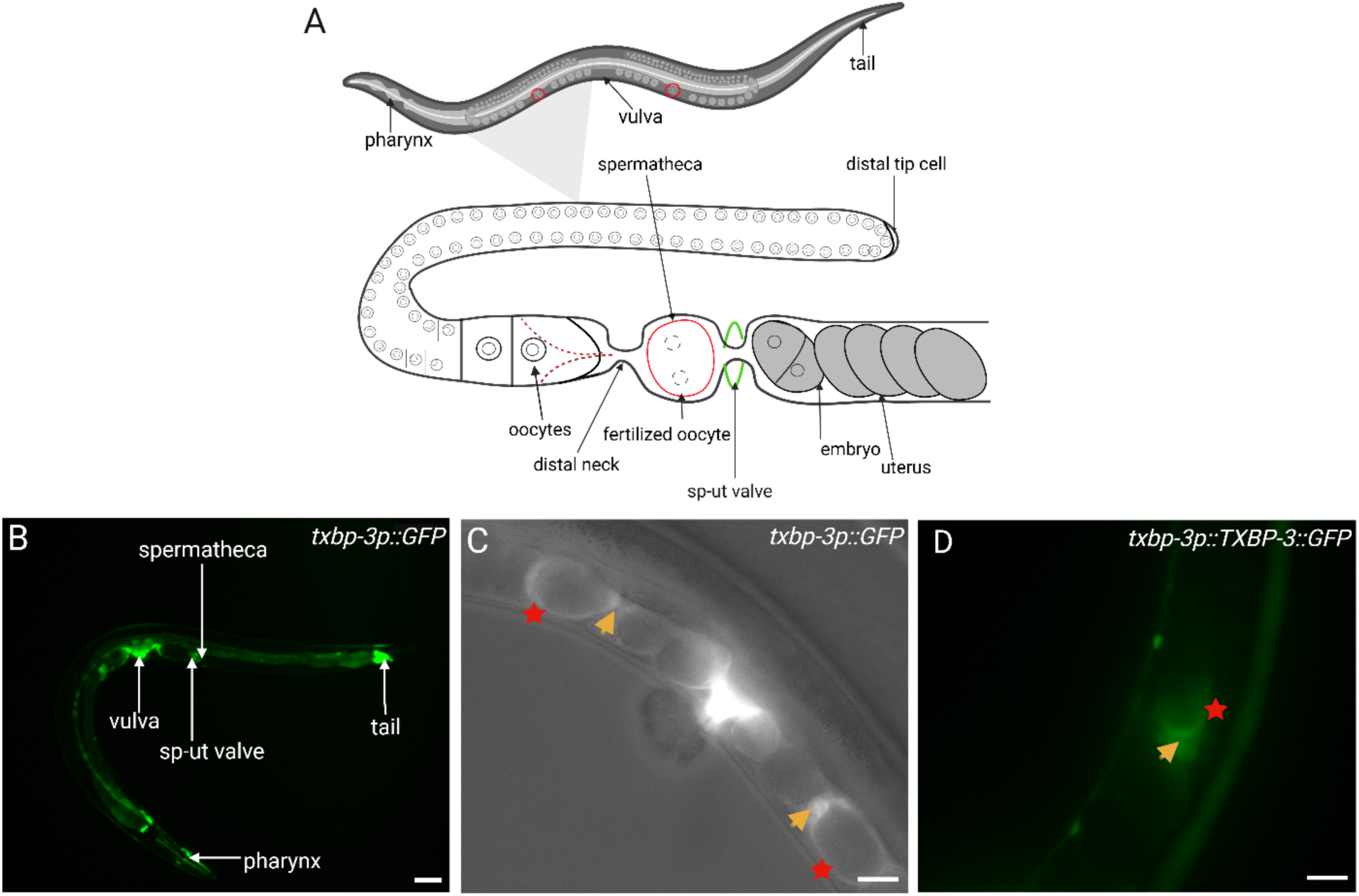
*txbp-3* is expressed in contractile tissues. (A) Schematic diagram of the *C. elegans* reproductive system showing the spermatheca and sp-ut valve connecting the gonad arm to the uterus. (B) Wide-field fluorescence image of a transgenic animal expressing *txbp-3p::GFP* transcriptional reporter. Expression is seen in the pharyngeal muscles, vulval muscles, spermatheca, sp-ut valve, and posterior intestine/tail (anterior left; ventral down). (C) DIC/GFP overlay image of the central region of the animal showing *txbp-3p::GFP* in the spermatheca (red star) and sp-ut valve (orange arrow) and (D) *txbp-3p::*TXBP-3::GFP. Scale bar = 20 µm. These images represent n > 20 animals. Fig. 1A was created with BioRender.com.

The signaling pathways that regulate spermathecal contractility resemble those in mammalian smooth and non-muscle cells (Sethi et al., 2017). Two parallel pathways control spermathecal contractility: the Ca^2+^-dependent pathway and the Rho-dependent pathway, both of which lead to the phosphorylation and activation of NMY-1/non-muscle myosin II (Bouffard et al., 2019). In the Ca^2+^-dependent pathway, the phospholipase PLC-1 cleaves phosphatidylinositol 4,5-bisphosphate (PIP_2_) into diacylglycerol (DAG) and inositol trisphosphate (IP_3_), triggering Ca^2+^ release from the endoplasmic reticulum through the IP₃ receptor ITR-1. Elevated Ca^2+^ levels activate the myosin light chain kinase MLCK-1, which phosphorylates myosin to initiate contraction (Kovacevic et al., 2013; Kelley et al., 2018). Upon oocyte entry, the mechanosensitive RhoGAP SPV-1/ARHGAP29 detaches from the membrane, enabling RHO-1 activation (Tan & Zaidel-Bar, 2015) by the PDZ-domain-containing RhoGEF RHGF-1/ARHGEF11 (Avivi Kela et al., 2023). Active RHO-1 then signals through LET-502/ROCK, which promotes myosin regulatory light chain phosphorylation both directly and by inhibiting myosin phosphatase MEL-11/MYPT (Somlyo & Somlyo, 2000). Together, these processes ensure coordinated spermathecal contraction and valve opening during ovulation, although their precise timing and spatial coordination remain incompletely understood.

PDZ-domain proteins (named for Postsynaptic density-95, Discs-large, and Zonula occludens-1) typically organize signaling complexes at the plasma membrane by enabling protein-protein interactions, supporting cellular integrity, and regulating processes such as cell adhesion, synaptic plasticity, and signal transduction, often under the control of post-translational modifications such as phosphorylation (Liu & Fuentes, 2018; Lee & Zheng, 2010).

TXBP-3 and its mammalian ortholog TAX1BP3/TIP-1 are atypical PDZ proteins consisting almost entirely of a single PDZ domain; rather than scaffolding complexes like most family members, they can bind target proteins and competitively displace them from their normal partners. For example, in mesenchymal progenitor cells, TAX1BP3 inhibits canonical Wnt/β-catenin and BMP/Smad signaling to reciprocally regulate osteogenic and adipogenic differentiation (Wang et al., 2023). TAX1BP3/TIP-1 has also emerged as a therapeutic target, where anti-TIP-1 antibodies enhance the efficacy of radiation therapy in non-small-cell lung cancer and glioblastoma by reducing proliferation and promoting cell death (Singh et al., 2023). Furthermore, measuring TAX1BP3 levels in melanoma can help predict whether a tumor will respond to immunotherapy (Aung et al., 2024). In zebrafish, TIP-1 participates in optic nerve regeneration, where its upregulation is linked to RhoA inactivation downstream of Wnt5b signaling (Matsukawa et al., 2017).

TAX1BP3 also regulates Ca^2+^ dynamics and contractility. In vascular smooth muscle cells, TAX1BP3 moves into the nucleus and binds to the transcriptional regulator YAP. By blocking YAP from interacting with its partner TEAD1, TAX1BP3 prevents excessive cell division and preserves the contractile state of the tissue. When TAX1BP3 is absent, these cells rapidly multiply and lose their contractility, resulting in abnormal blood vessel thickening (Yang et al., 2025). In cardiomyocytes, TAX1BP3 supports Ca^2+^ homeostasis, and its loss derepresses the TRPV4 channel, causing TRPV4-mediated Ca^2+^ leak and arrhythmogenic cardiomyopathy (Perelli et al., 2025).

In this study, we identify *C. elegans* TXBP-3 as a dosage-dependent regulator of spermathecal contractility. We show that *txbp-3* is expressed in multiple contractile tissues, including the pharynx, vulval muscles, spermatheca, and the sp-ut valve. Using CRISPR/Cas9-mediated deletion and tissue-specific overexpression, we demonstrate that TXBP-3 fine-tunes spermathecal contractility, with both its loss and overexpression disrupting normal transit. TXBP-3 promotes the activation of UNC-43, the *C. elegans* CaMKII ortholog. Furthermore, TXBP-3 functions in parallel with the RhoGAP SPV-1 to regulate contractile actomyosin organization in the spermatheca and maintain embryo shape and structural integrity.

## Results

### TXBP-3 is expressed in contractile tissues of *C. elegans*

To investigate the role of TXBP-3 in *C. elegans*, we analyzed its expression pattern using both transcriptional *txbp-3p::GFP* and translational *txbp-3p::TXBP-3::GFP* reporter lines. Both reporters showed that TXBP-3 is widely expressed in multiple contractile tissues, including the somatic gonad and the vulval muscles (Fig. 1B). Within the gonad, TXBP-3 is expressed in the spermatheca and the sp-ut valve (Fig. 1C). The TXBP-3::GFP fusion protein is distributed in the cytoplasm of the spermathecal cells and the sp-ut valve (Fig. 1D). This expression pattern suggests that TXBP-3 may regulate oocyte transit and valve function.

### TXBP-3 is a highly conserved PDZ domain protein

The *C. elegans txbp-3* gene is predicted to encode a 124-amino-acid protein composed of a single PDZ domain. Phylogenetic analysis indicates that *txbp-3* is highly conserved across Metazoa, with clear orthologs in other animals (Fig. 2A). AlphaFold modeling predicts that TXBP-3 adopts a β-sheet PDZ core, which is structurally similar across orthologs, with short divergent flexible loop regions (Jumper et al., 2021) (Fig. 2B). Multiple sequence alignment indicates that amino acid conservation is concentrated within this structural core (Fig. 2C).

**Figure 2.**
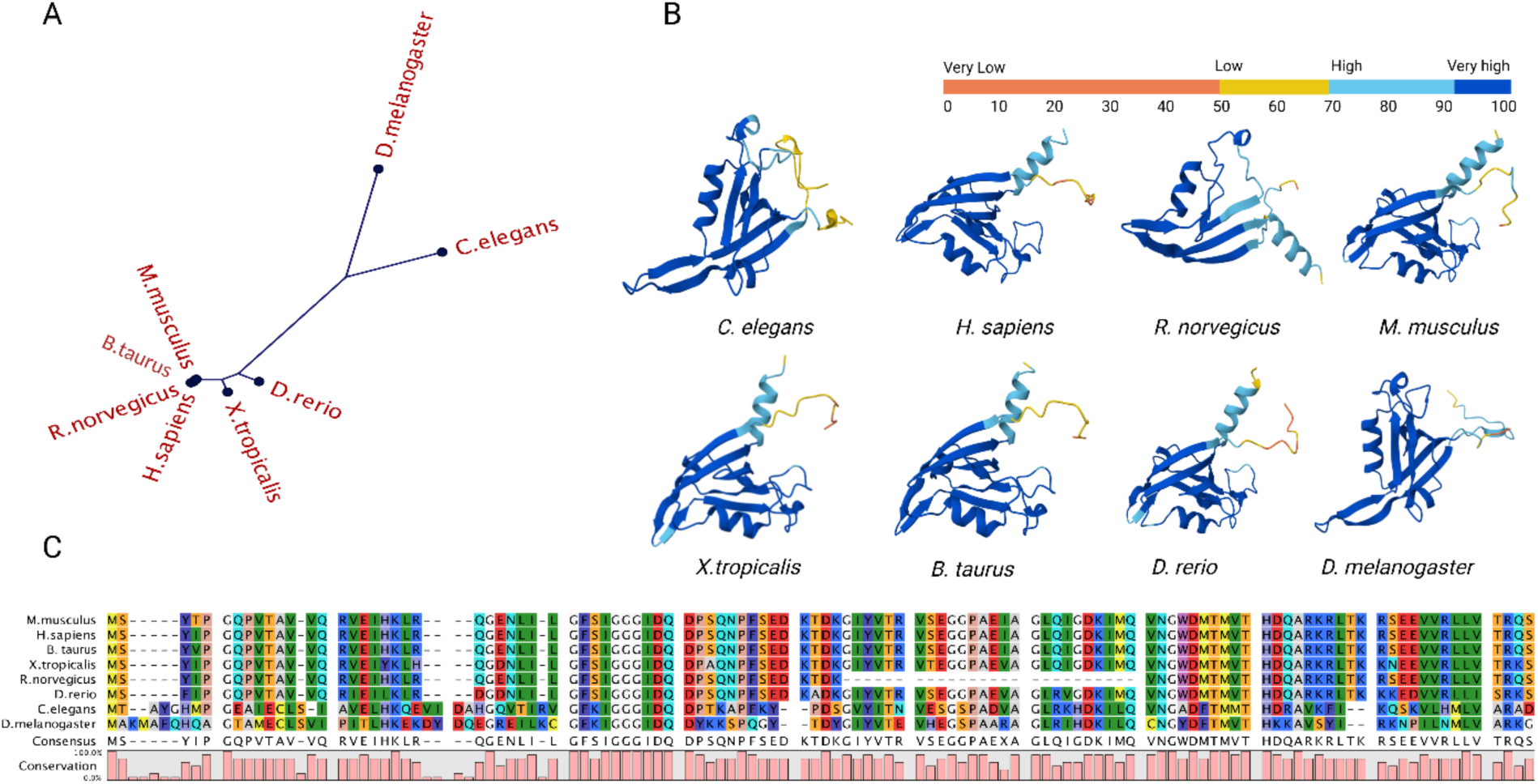
Structural and sequence conservation of TXBP-3. (A) Unrooted phylogenetic tree of TXBP-3 orthologs across Metazoa. (B) AlphaFold-predicted tertiary structures of TXBP-3 (*C. elegans* ID: A8Y3F3); orthologs correspond to TAX1BP3 in *H. sapiens*, *R. norvegicus*, *M. musculus*, *D. rerio*, *B. taurus, X. tropicalis*; and CG3402 in *D. melanogaster*. Structures are colored by model confidence (pLDDT): dark blue, very high (>90); light blue, high (70–90); yellow, low (50–70); orange, very low (<50). (C) Multiple sequence alignment of TXBP-3 orthologs. The pink histogram indicates conservation at each residue, highlighting identity within the core binding groove.

As seen in the phylogenetic tree, AlphaFold models, and sequence alignment, TXBP-3 is more closely related to the *D. melanogaster* ortholog than the mammalian proteins.

### TXBP-3 is a dosage-dependent regulator of oocyte transit

To investigate the function of TXBP-3, we utilized CRISPR/Cas9-mediated genome editing to create *txbp-3(knu748),* a 1005-bp deletion allele that removes the entire coding sequence and a portion of the promoter. We confirmed the deletion through PCR and sequencing. The resulting null mutant *txbp-3(Δ)* was backcrossed four times to N2.

Oocyte transit through the spermatheca occurs in a coordinated manner consisting of three phases: entry (when the oocyte enters the spermatheca), dwell (when the oocyte remains completely enclosed within the spermatheca), and exit (when the embryo leaves the spermatheca and enters the uterus) (Fig. 3A; Movie S1). For a successful transit, the distal bag must contract to push the embryo out through the sp-ut valve. Therefore, transit times depend on the relationships between neck, bag, and valve contractility (Fig. S1). We next characterized the *txbp-3(Δ)* spermathecal transits by DIC time-lapse microscopy (Movie S2), and found no significant differences in entry, dwell, or exit times in the *txbp-3(Δ)* mutants (Fig. 3B-D). Most ovulations were successful (9/10), with one instance where the distal neck closed prematurely, severing the oocyte and resulting in retention of a small fragment in the gonad arm (Table S1).

**Figure 3.**
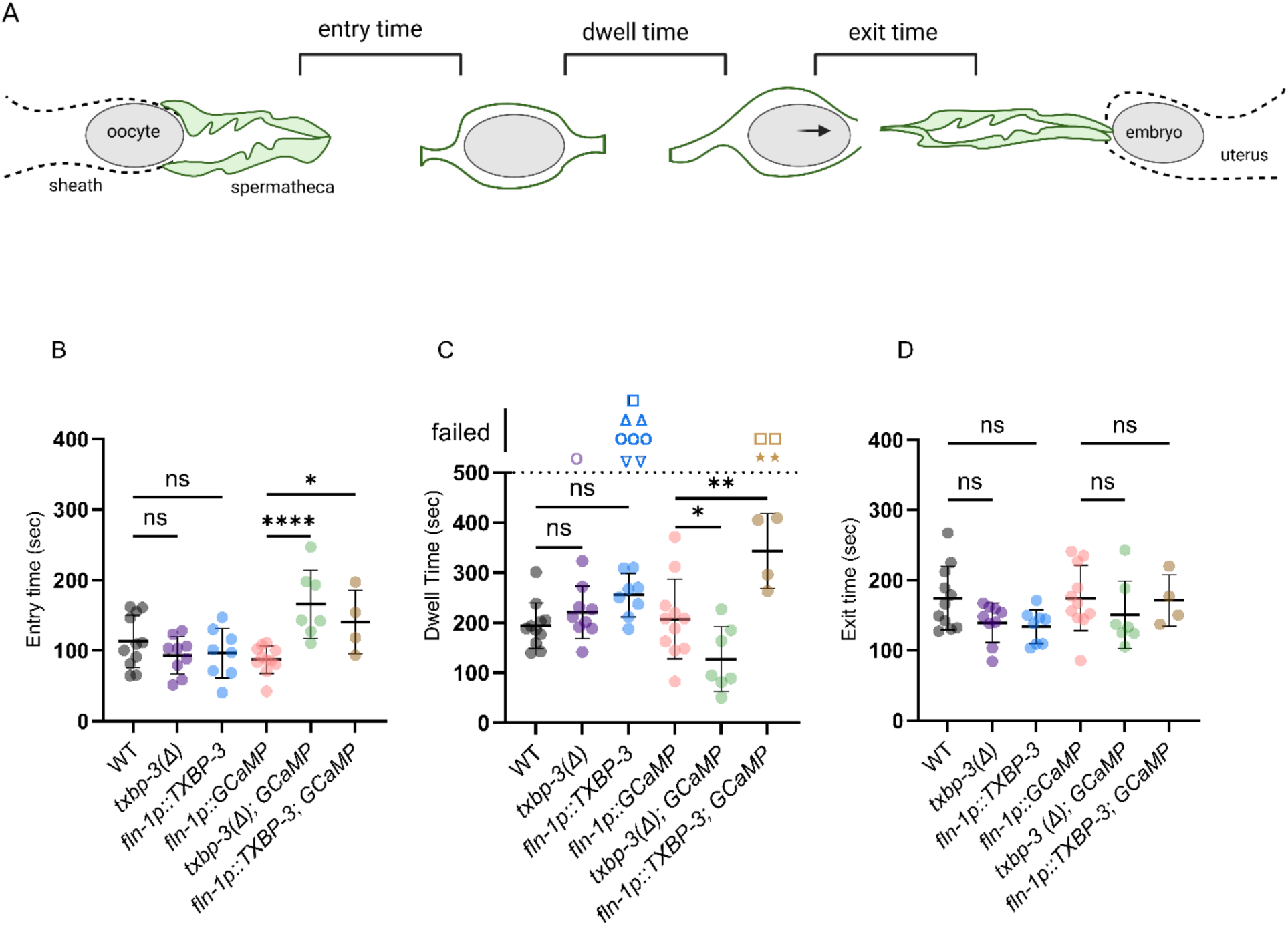
TXBP-3 regulates oocyte transit in a dosage-dependent manner. (A) Schematic of oocyte transit defining the three coordinated phases of ovulation: entry time (neck opening to closure), dwell time (neck closure to sp-ut valve opening), and exit time (sp-ut opening until the embryo enters the uterus). (B-D) Quantification of (B) entry, (C) dwell, and (D) exit times across indicated genotypes in both N2 and GCaMP3 backgrounds. In (C), symbols above the 500 sec dashed line denote transit failures: severing due to tight neck (circles), severing due to tight sp-ut valve (inverted triangle), trapping (squares), pushback (star) and rupture (triangles). Data are presented as mean ± SD. Statistical significance was determined using an ordinary one-way ANOVA followed by Šídák’s multiple comparisons test (\**p* < 0.05, \*\**p* < 0.01, \*\*\**p* < 0.001, \*\*\*\**p* < 0.0001, ns = not significant).

Because transits can be difficult to visualize in N2 animals, we repeated the experiments utilizing the genetically encoded fluorescent calcium sensor GCaMP3. In this background, *txbp-3(Δ)* resulted in significantly slower entry times (Fig. 3B) and shorter dwell times (Fig. 3C) than control *fln-1p::GCaMP* animals. This may be because GCaMP3 acts as a Ca^2+^ buffer, sensitizing the tissue to *txbp-3* loss. Both slow entry and shorter dwell times can result from reduced contractility. Slower entry occurs because the less-contractile spermathecal neck cells are delayed in closing around the oocyte (Kovacevic et al., 2013), and a shorter dwell time can result from a less-contractile sp-ut valve opening more easily (Kelley et al., 2020).

Because the spermatheca in the *txbp-3(Δ)* animals seemed to be less contractile, we hypothesized that overexpression might increase spermathecal contractility. We overexpressed TXBP-3 in the spermatheca using the *fln-1* promoter (*fln-1p::*TXBP-3) on a multi-copy array (Kovacevic & Cram, 2010). Overexpression of TXBP-3 disrupted transits (Movie S3), resulting in transit failures in both backgrounds and significantly longer dwell times in the sensitized GCaMP3 background compared to *fln-1p::GCaMP* (Fig. 3C). Entry times were also significantly slower in *fln-1p::*TXBP-3*; GCaMP* animals compared to *fln-1p::GCaMP* (Fig. 3B). The slower entry likely reflects a hypercontracted neck that resists oocyte entry, whereas the prolonged dwell time likely reflects a hypercontracted sp-ut valve.

In the *fln-1p::*TXBP-3 overexpression line, the overall transit success rate was 61.5% (8/13). In the remaining transits, the spermathecal neck cells closed prematurely around the oocyte and severed it (23.1%, 3/13), the sp-ut valve closed and severed the oocyte (15.4%, 2/13), the oocyte ruptured (15.4%, 2/13), or the embryo failed to exit (7.7%, 1/13) and remained in the spermatheca indefinitely. Some transits displayed multiple defects (Table S1).

Similarly, transit failures were observed in the sensitized background (*fln-1p::*TXBP-3*; GCaMP*), which had a failure rate of 50% (4/8). Transit failures in this line included failure of oocyte exit (25%, 2/8) and two in which the oocyte entered the spermatheca but was immediately pushed back into the gonad arm (25%, 2/8) (Table S1).

### TXBP-3 overexpression leads to transit failures and disrupted calcium signaling in the spermatheca

Actomyosin contractility in the spermatheca is, in part, regulated by Ca^2+^ release from internal stores, which activates MLCK-1/myosin light chain kinase (Kelley et al., 2018). To examine if TXBP-3 affects Ca^2+^ release during oocyte transit, we measured Ca^2+^ using the GCaMP3 sensor by capturing time-lapse movies of GCaMP signal at one frame per second (Movies S4-S6). To visualize the spatiotemporal aspects of this Ca^2+^ signaling, we plotted the data both as 1D traces of total Ca^2+^ levels (Fig. 4A-C) and as 2D kymograms with time on the x-axis and space on the y-axis (Fig. 4A’-C’).

**Figure 4.**
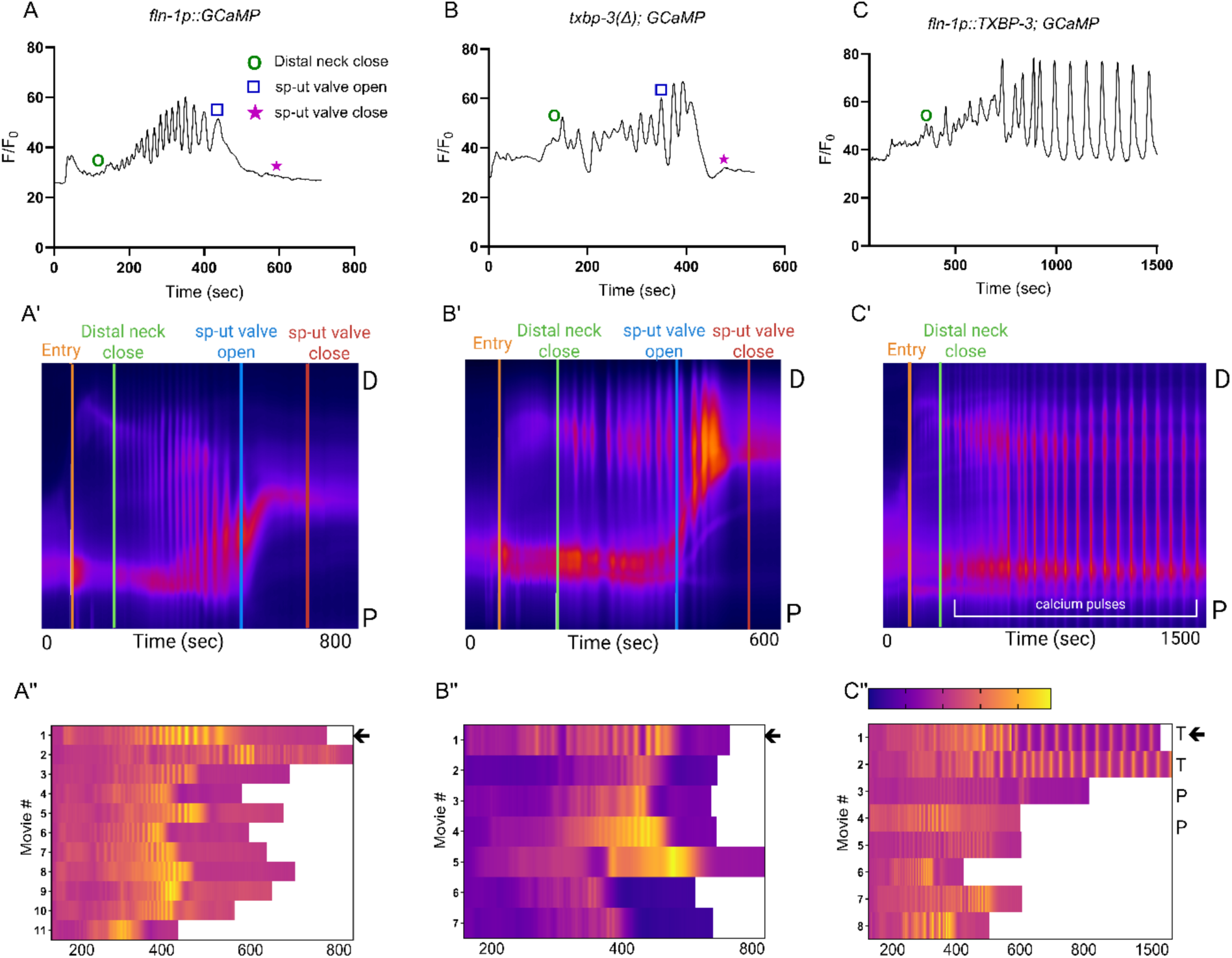
TXBP-3 regulates Ca^2+^ signaling and prevents mechanical transit failure. (A-C) Normalized 1D Ca^2+^ traces, (A’-C’) 2D spatiotemporal kymograms, and (A”-C”) heatmaps for (A) WT, (B) *txbp-3(Δ)*, and (C) *fln-1p::*TXBP-3 animals (all expressing *fln-1p::GCaMP*). In (A-C), symbols denote distal neck closure (green circle), sp-ut valve opening (blue square), and valve closure (pink star). In (A′-C′), the y-axis shows position along the spermatheca, from distal (D, top) to proximal (P, bottom). The bracket in (C’) highlights sustained Ca^2+^ during a trapped oocyte transit. In (A”-C”), warmer colors indicate higher Ca^2+^ intensity; arrows mark the representative transits shown in A-C (T: trapped, P: pushed back).

In WT animals (*fln-1p::GCaMP*), oocyte entry triggers a Ca^2+^ wave that starts at the distal neck and propagates rhythmically toward the sp-ut valve. This steady propagation is visible in representative traces and kymograms (Fig. 4A, A’, A’’; Movie S4). The *txbp-3(Δ)* animals exhibit a similar rhythmic pattern (Fig. 4B, B’; Movie S5); however, the overall Ca^2+^ signal intensity is weaker and shows increased variability (Fig. 4B’’, Fig. S2).

TXBP-3 overexpression (*fln-1p::*TXBP-3) can lead to two different outcomes. When the embryo exits successfully, Ca^2+^ transients are similar to WT (Fig. S2). However, when oocytes fail to exit, strong and continuous Ca^2+^ pulses are observed (Fig. 4C, C’; Movie S6). These pulses are insufficient to expel the embryo into the uterus, likely due to excess constriction of the sp-ut valve. Supporting this assertion, in two instances, the embryo was pushed back through the entry neck and into the gonad arm (indicated by “P” in Fig. 4C’’).

Taken together, these results suggest TXBP-3 regulates spermathecal contractility and oocyte transit, with a potential role in Ca^2+^ signal coordination.

### TXBP-3 promotes UNC-43 activation in the spermatheca

Earlier high-throughput studies suggested that TXBP-3 physically interacts with UNC-43, the *C. elegans* ortholog of CaMKII (Lenfant et al., 2010). The UNC-43 C-terminus ends in a sequence matching the PDZ-binding consensus (Nourry et al., 2003) (Fig. 5A). AlphaFold structural predictions support this potential interaction, suggesting that the UNC-43 C-terminus might interface with the α-helix of the TXBP-3 PDZ domain (Evans et al., 2021) (Fig. 5B).

**Figure 5.**
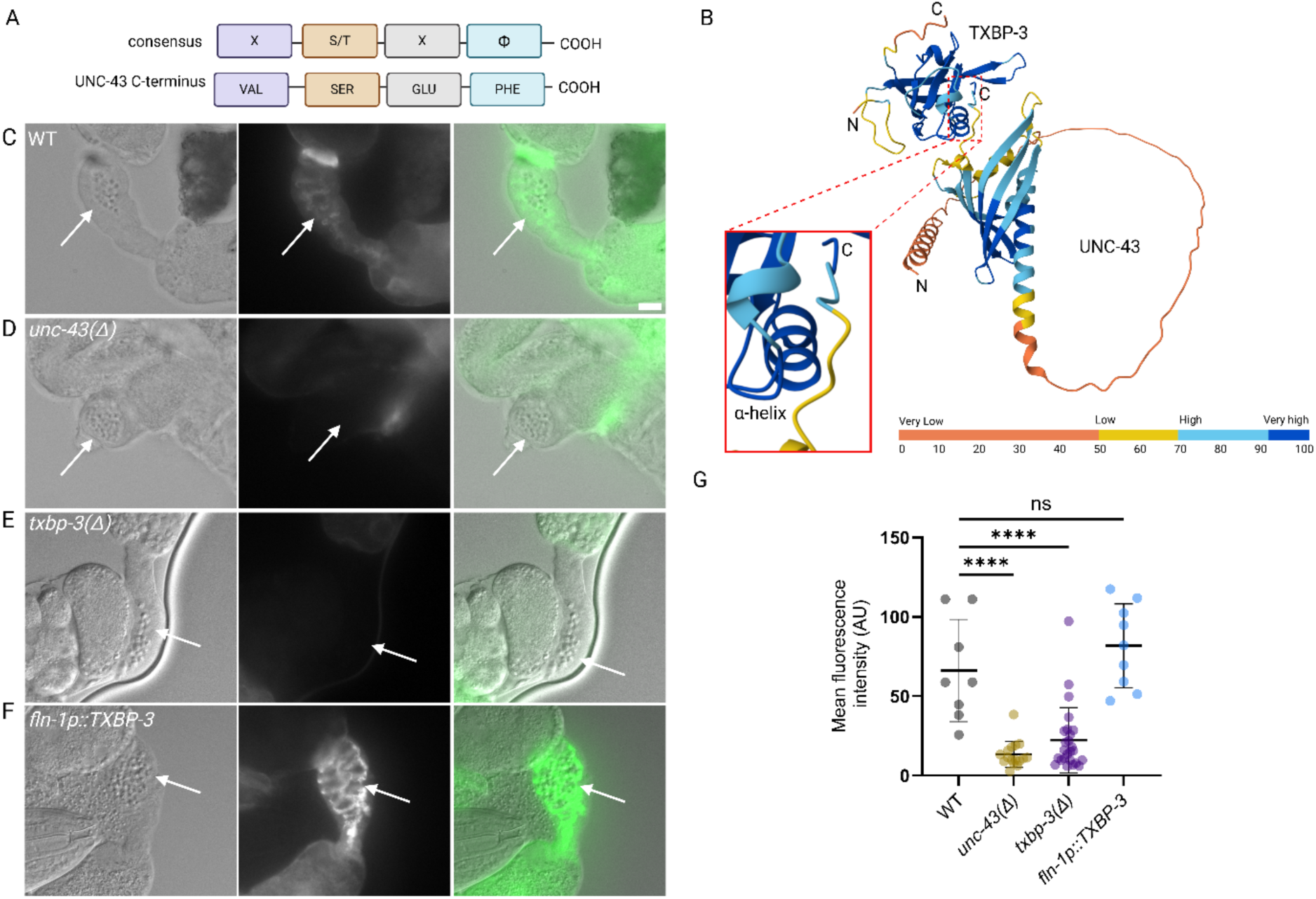
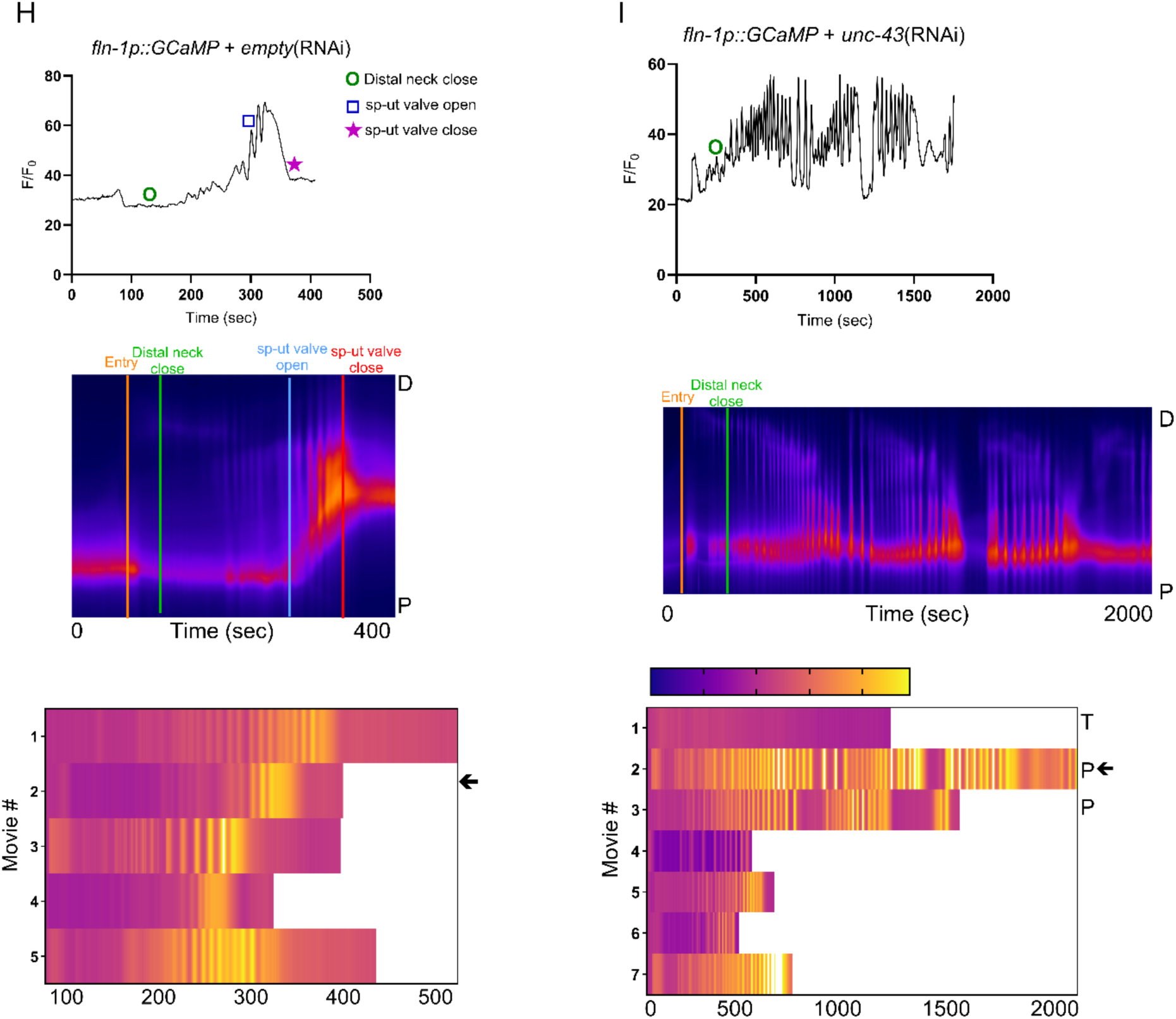
TXBP-3 promotes UNC-43 activation. (A) Alignment of the UNC-43 C-terminus with a PDZ-binding motif. (B) AlphaFold-Multimer structural prediction of the TXBP-3/UNC-43 complex, colored by model confidence (pLDDT). (C-F) Representative immunofluorescence images (DIC, phospho-UNC-43, and merged channels) of the dissected spermatheca (white arrows) in (C) WT, (D) *unc-43(Δ)*, (E) *txbp-3(Δ)*, and (F) *fln-1p::*TXBP-3 animals. Scale bar: 10 µm. (G) Quantification of phospho-UNC-43 mean fluorescence intensity (MFI). Data are presented as mean ± SD; one-way ANOVA followed by Dunnett’s multiple comparisons test (\*\*\*\**p* < 0.0001, ns = not significant). (H-I) Representative Ca^2+^ traces, spatiotemporal kymograms, and transit heatmaps in *fln-1p::GCaMP* animals treated with (H) *empty*(RNAi) or (I) *unc-43*(RNAi). Arrows mark the transits depicted in the kymograms above (T: trapped, P: pushed back).

To determine if TXBP-3 regulates UNC-43 activity, we performed immunofluorescence on dissected gonads using a CaMKII-specific monoclonal antibody that binds phospho-Thr286 and indicates UNC-43 activity. In WT animals, phosphorylated UNC-43 is observed faintly across the spermatheca and more brightly in the sp-ut valve (Fig. 5C, G). Importantly, this fluorescent signal was absent in the *unc-43(js125)* deletion mutant, confirming antibody specificity (Fig. 5D, G). We observed lower signal levels in *txbp-3(Δ)* mutants (Fig. 5E, G). With *fln-1p::*TXBP-3 overexpression, we observed similar levels of phosphorylated UNC-43 signal to WT in the spermatheca and sp-ut valve (Fig. 5F, G). Lower signal in the *txbp-3(Δ)* mutants suggests that TXBP-3 stimulates UNC-43 activity in the spermatheca.

Next, we asked whether RNAi-mediated depletion of *unc-43* results in defects in Ca^2+^ signaling. Control WT (*fln-1p::GCaMP*) animals treated with *empty*(RNAi) showed normal rhythmic Ca^2+^ waves and successful transits (Fig. 5H; Movie S7). Depletion of *unc-43* in the WT GCaMP background results in a variety of defects, including trapping and abnormal movement of the oocyte in the spermatheca (Fig. 5I; Movie S8). In these instances, spatiotemporal aspects of Ca^2+^ were also disrupted. Despite these defects, *unc-43* depletion does not alter the overall transit parameters in the GCaMP background, possibly because the GCaMP sensor itself is buffering the Ca^2+^ availability and masking the phenotype (Fig. S3). When *unc-43* was depleted in the *fln-1p::*TXBP-3; GCaMP background, we observed no significant differences in transit times (Fig. S3) or Ca^2+^ metrics (Fig. S4) compared to *empty*(RNAi) controls, perhaps because the HT115(DE3) bacterial strain used for feeding RNAi suppresses the hypercontractile phenotype observed in animals expressing *fln-1p::*TXBP-3.

### TXBP-3-mediated hypercontractility is independent of the RhoGEF RHGF-1

In addition to Ca^2+^-activated regulation of MLCK-1, spermathecal contractility depends on the activation of the small GTPase RHO-1 (Kelley et al., 2020). Given the observed hypercontractile phenotypes, we suspected that TXBP-3 might interact with the Rho signaling pathway. Rho is activated by guanine nucleotide exchange factors (GEFs) that increase GTP-bound Rho. Previous studies identified three GEFs as regulators of spermathecal transit: RHGF-1, UNC-73, and Y105E8A.25 (Avivi Kela et al., 2023). As RHGF-1 contains a PDZ domain, we hypothesized that RHGF-1 might interact with a single-PDZ-domain protein like TXBP-3.

To investigate a possible functional relationship between TXBP-3 and RHGF-1, we analyzed genetic interactions between *txbp-3(Δ)* and *rhgf-1(ok880)*. In both the single *rhgf-1(Δ)* and *txbp-3(Δ); rhgf-1(Δ)* double mutants, oocyte entry time and dwell time were not significantly different from WT (Fig. 6A-B). As expected (Avivi Kela et al., 2023), the single mutant *rhgf-1(Δ)* showed significantly faster exit due to reduced contractility in the sp-ut valve (Fig. 6C).

**Figure 6.**
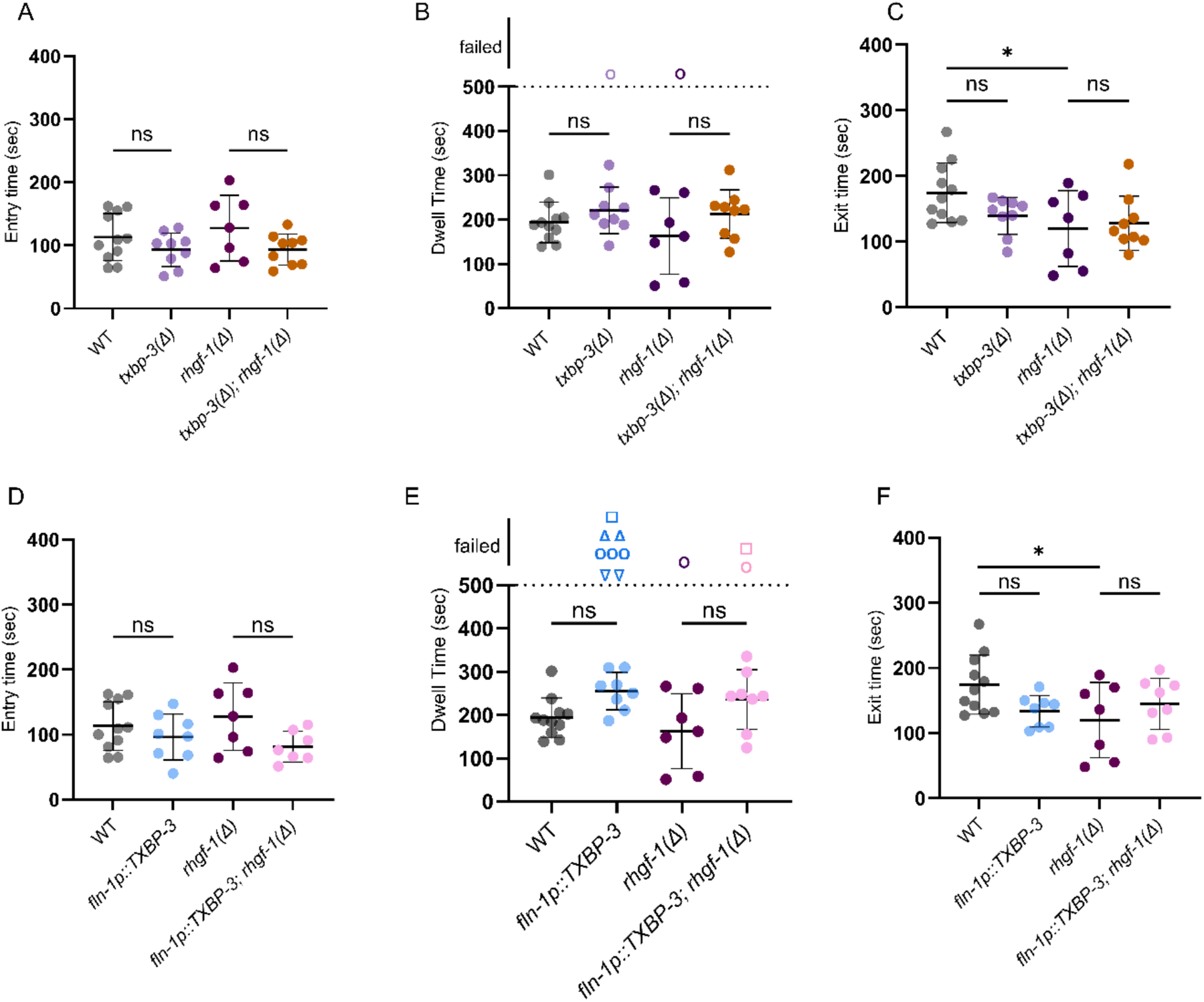
TXBP-3-driven contractility is independent of RHGF-1. (A-C) Oocyte transits in loss-of-function mutants. Scatter plots quantifying (A) entry time, (B) dwell time, and (C) exit time in WT, *txbp-3(Δ)*, *rhgf-1(Δ),* and *txbp-3(Δ)*; *rhgf-1(Δ)* double mutants. (D-F) Oocyte transit in overexpression backgrounds. Scatter plots quantifying (D) entry time, (E) dwell time, and (F) exit time across indicated genotypes. In (B, E), symbols plotted above 500 sec indicate transit failures, including oocyte severing due to tight neck (circles), trapping (squares), oocyte severing due to tight sp-ut valve (inverted triangle), and rupture (triangles). Data are presented as mean ± SD. Statistical significance was determined using a one-way ANOVA followed by Šídák’s multiple comparisons test (\**p* < 0.05, ns = not significant).

Because RHGF-1 stimulates contractility by activating RHO-1, we predicted that the loss of *rhgf-1* might reduce the increased contractility seen in *fln-1p::*TXBP-3 animals. In *fln-1p::*TXBP-3 animals, increased valve contractility leads to prolonged dwell times and frequent transit failures (Fig. 6E). Yet, in the *fln-1p::*TXBP-3*; rhgf-1(Δ)* animals, entry and exit times remained comparable to the *fln-1p::*TXBP-3 overexpression line (Fig. 6D, F), and dwell times remained similarly elevated (Fig. 6E). This suggests that TXBP-3-mediated hypercontraction does not require RHGF-1, and that the loss of RHGF-1 is not sufficient to lessen the effects of TXBP-3 overexpression.

### TXBP-3 acts in parallel with SPV-1 to regulate spermathecal contractility

Rho is negatively regulated by GTPase-activating proteins (GAPs). The GAP SPV-1 inhibits RHO-1, and without SPV-1, the spermatheca is hypercontractile (Tan & Zaidel-Bar, 2015). To investigate the functional relationship between TXBP-3 and SPV-1, we analyzed genetic interactions between *txbp-3* and *spv-1* mutants (Fig. 7A-C). Transit times were faster in all *spv-1(ok1498)* backgrounds, resulting in significantly shorter entry and exit times compared to WT (Fig. 7A, C). The greater spermathecal contractility in these mutants propels the oocyte through the spermatheca more rapidly.

**Figure 7.**
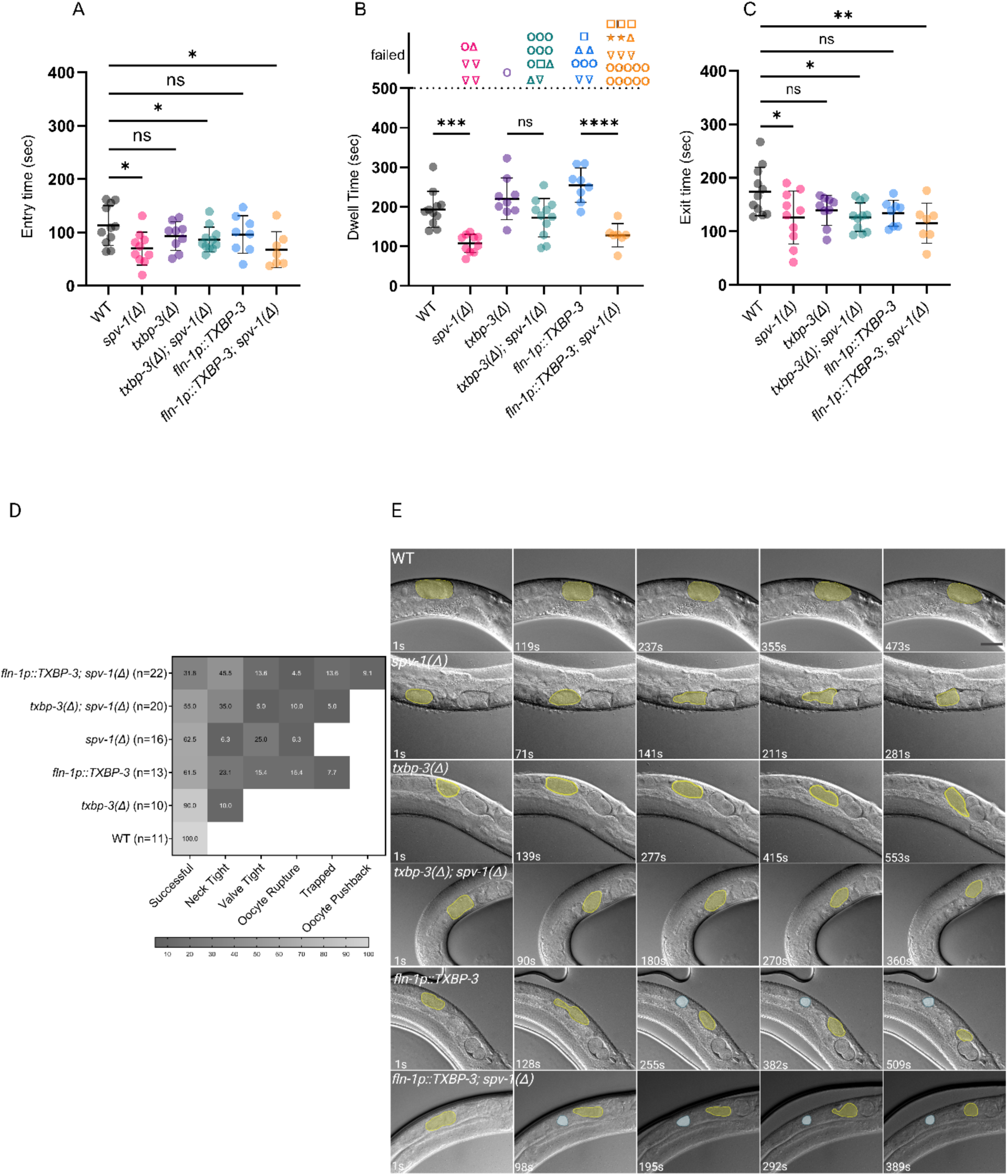
TXBP-3 and SPV-1 act in parallel to regulate spermathecal contractility. (A-C) Quantification of oocyte (A) entry, (B) dwell, and (C) exit times across genotypes. In (B), symbols denote specific transit outcomes: successful transits (solid circles, <500 sec) and mechanical failures (>500 sec), including severing due to tight neck (open circles), trapping (squares), severing due to tight sp-ut valve (inverted triangle), pushback (star), and rupture (triangles). (D) Heat map of oocyte transit outcomes across genotypes; color intensity and values reflect % of animals (of n) per outcome, which are not mutually exclusive (see Table S1). (E) Representative DIC time-lapse images of oocyte transit stages. Yellow outlines highlight the main oocyte body; blue shading depicts severed oocyte fragments. Scale bar: 20 µm. Statistical significance was determined using a one-way ANOVA. Entry and exit times (A, C) were analyzed with Dunnett’s multiple comparisons test; dwell time (B) was analyzed with Tukey’s multiple comparisons test (\**p* < 0.05, \*\**p* < 0.01, \*\*\**p* < 0.001, \*\*\*\**p* < 0.0001, ns = not significant).

While both *spv-1(Δ)* and *fln-1p::*TXBP-3 cause hypercontractility, their effects differ. Loss of SPV-1 triggers RHO-1-driven hypercontractility in the distal spermatheca and in the sp-ut valve. The distal spermatheca contracts strongly from the rear of the bag to force the oocyte through a hypercontracted valve. In contrast, TXBP-3 overexpression drives hypercontractility at both the distal neck and the sp-ut valve, causing premature neck closure and a valve that acts as a closed sphincter that delays or blocks oocyte exit. In the *spv-1(Δ)* mutants (Movie S9), oocyte transits were disrupted due to valve tightness (25%, 4/16; Table S1), where the sp-ut valve closed prematurely and severed the oocytes, resulting in the formation of pear-shaped embryos. In one instance, the distal neck closed prematurely, severing the oocyte and leaving a small fragment in the gonad arm.

We next tested if TXBP-3 is required for this hypercontractility. In the *txbp-3(Δ); spv-1(Δ)* double mutants (Movie S10), dwell times were not statistically different from *txbp-3(Δ)* single mutants (Fig. 7B). This suggests that the loss of TXBP-3 can balance out the loss of SPV-1, resulting in partially normalized spermathecal bag contractility. However, overall transit success rate dropped to 55% (11/20). Mechanical failures occurred in the remaining transits, including premature closure of the spermathecal neck, severing the oocyte (35%, 7/20), severing by the sp-ut valve (5%, 1/20), oocyte rupture (10%, 2/20), and trapped oocytes that failed to exit (5%, 1/20), where the embryo remained in the spermatheca indefinitely (Fig. 7D, E; Table S1). This suggests premature activation of RHO-1 in the neck still occurs in the *txbp-3(Δ); spv-1(Δ)* double mutants.

As expected, in the *fln-1p::TXBP-3; spv-1(Δ)* animals (Movie S11), the overall successful transit rate was lowest (31.8%, 7/22), with the exit time significantly shorter than WT (Fig. 7C). Hypercontractility was observed at the spermathecal neck (45.5%, 10/22) and valve (13.6%, 3/22). Defects including oocyte rupture (4.5%, 1/22), trapped oocytes (13.6%, 3/22), and oocytes that were pushed back into the gonad arm (9.1%, 2/22) were also observed (Fig. 7D; Table S1). These data demonstrate that the simultaneous disruption of SPV-1 and overexpression of TXBP-3 leads to increased contractility and significant spermathecal transit defects.

### Uncoordinated spermathecal and sp-ut valve contractility compromises embryo morphology and structural integrity

To determine whether defects in spermathecal contraction result in altered embryo shapes, we measured the area and aspect ratio of the embryos (Fig. 8A-F). In WT and *txbp-3(Δ),* embryos were consistent, with a similar area (∼1300 µm^2^) and aspect ratio (∼1.5), with no fragmentation (Fig. 8A, A’, B, B’, G). These findings suggest that the loss of *txbp-3* alone does not influence embryo shape.

**Figure 8.**
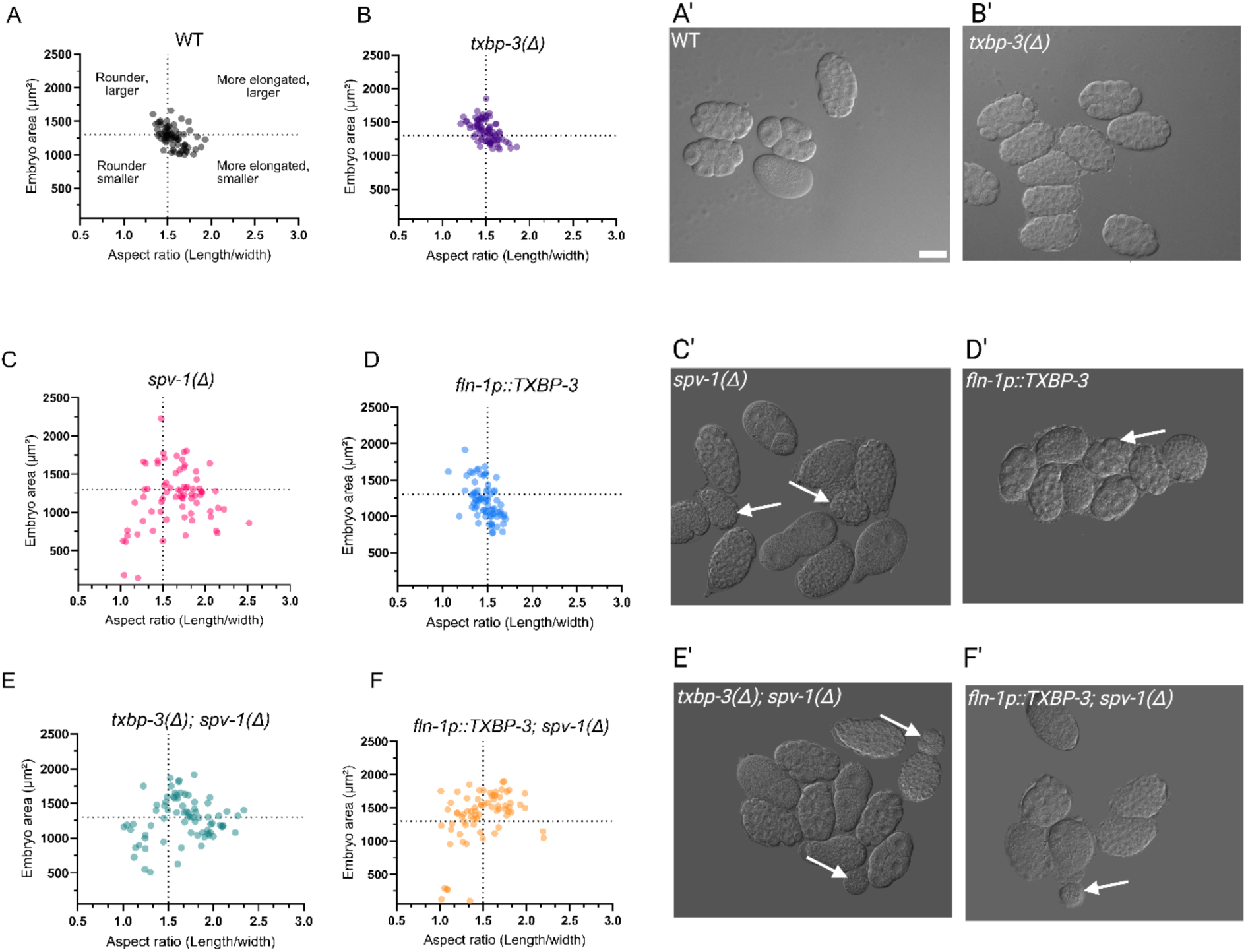
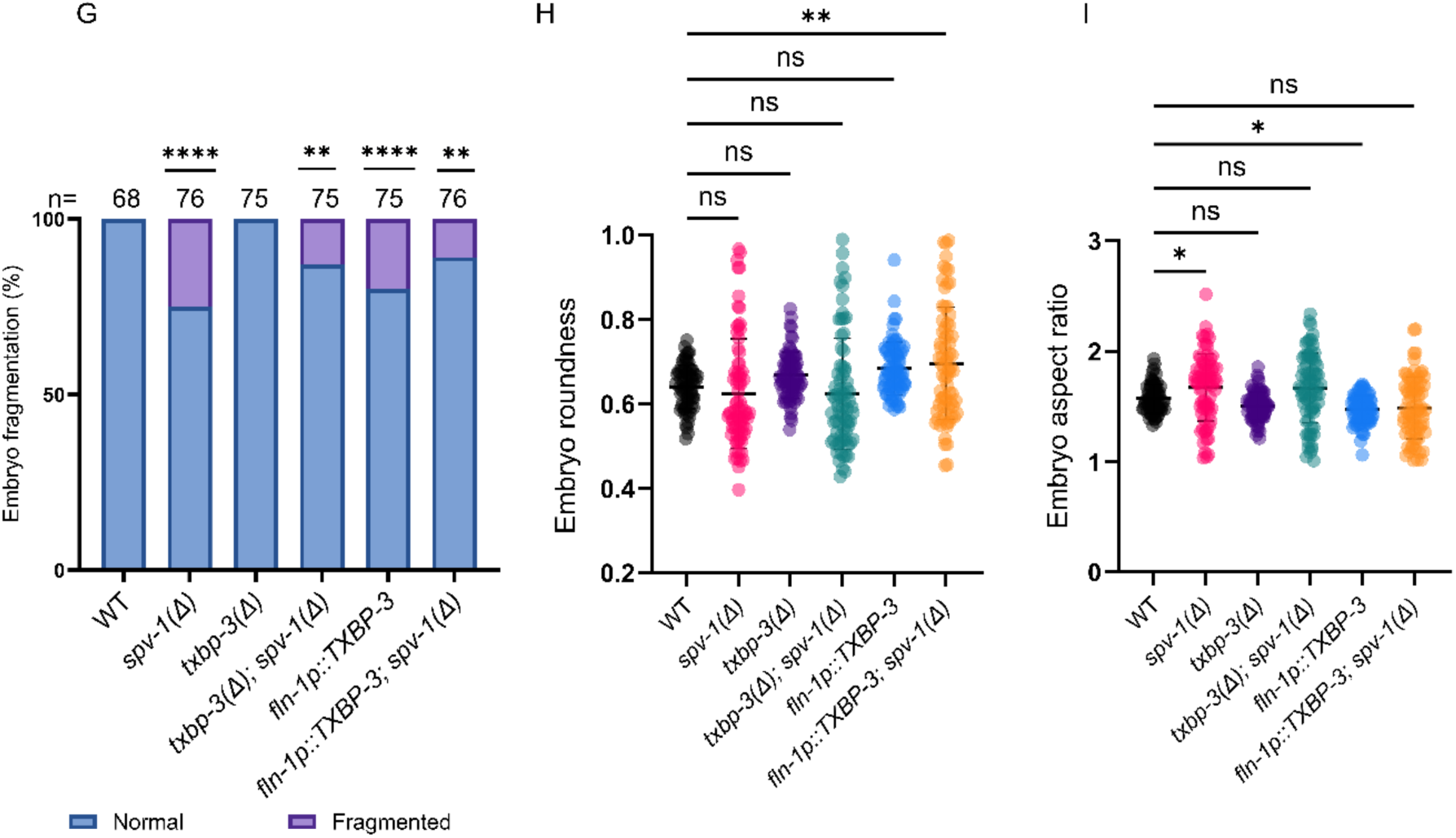
Uncoordinated contractility of the spermatheca and sp-ut valve disrupts embryo structural integrity. (A-F) Scatter plots show embryo area (µm^2^) versus the aspect ratio (length/width) for (A) WT, (B) *txbp-3(Δ),* (C) *spv-1(Δ)*, (D) *fln-1p::*TXBP-3, (E) *txbp-3(Δ); spv-1(Δ)*, and (F) *fln-1p::*TXBP-3*; spv-1(Δ)*. Dashed lines indicate WT mean area and aspect ratio. (A’-F’) Representative DIC images of embryos corresponding to genotypes in (A-F). Arrows indicate embryo fragments. Scale bar = 20 µm. (G) Stacked bar chart showing the percentage of normal (blue) versus fragmented (purple) embryos; brackets show comparisons with WT. Sample sizes (n) are indicated above each bar. (H-I) Quantification of embryo (H) roundness and (I) aspect ratio across genotypes. Data are mean ± SD. Statistical significance was determined by Fisher’s exact test for categorical frequencies (G) and one-way ANOVA with Dunnett’s multiple comparisons test for continuous variables (H, I) (\**p* < 0.05, \*\**p* < 0.01, \*\*\*\**p* < 0.0001, ns = not significant).

In contrast, *fln-1p::TXBP-3* and *spv-1(Δ)* animals with hypercontractile valves or necks exhibited embryo fragmentation (Fig. 8G). The *spv-1(Δ)* mutants showed the highest fragmentation rate at 25% (19/76) (Table S2). Without SPV-1 to relax the valve and neck, the oocytes become pinched, resulting in a significantly increased aspect ratio (Fig. 8C, C’, I). These results are consistent with previous reports (Tan & Zaidel-Bar, 2015; Bouffard et al., 2019). The hypercontractility of the *fln-1p::TXBP-3* overexpression line also resulted in embryo fragmentation (20%, 15/75; Fig. 8D’,G; Table S2). In contrast to the elongated *spv-1(Δ)* embryos, *fln-1p::*TXBP-3 embryos were rounder and exhibited a significantly lower aspect ratio (Fig. 8D, H, I).

To further investigate whether the genetic interaction of *spv-1(Δ)* and *txbp-3* affects embryo shape, we created double mutants with *spv-1(Δ)* in the *txbp-3(Δ)* background. As expected, the *txbp-3(Δ); spv-1(Δ)* double mutant exhibited lower fragmentation (13.3%, 10/75; Table S2) than the *spv-1* single mutant, again suggesting that loss of TXBP-3 can partially compensate for loss of SPV-1 (Fig. 8G).

Based on the hypercontractile phenotypes of the single mutants, we hypothesized that the combined loss of Rho inhibition (via *spv-1*) and increased TXBP-3 activity would exacerbate the phenotype, resulting in higher rates of embryo fragmentation. Surprisingly, the *fln-1p::*TXBP-3*; spv-1(Δ)* animals exhibited lower fragmentation frequencies (10.5%, 8/76; Table S2) than the *spv-1* single mutant (Fig. 8G). This relative reduction in fragmentation is likely due to entry defects such as severing by neck closure and oocyte reflux into the gonad arm, which preclude the observation of sp-ut valve phenotypes.

While the *txbp-3(Δ); spv-1(Δ)* double mutant embryos are similar to *spv-1(Δ)* single mutants in roundness and aspect ratio (Fig. 8E, E’, H, I), the *fln-1p::*TXBP-3*; spv-1(Δ)* embryos were rounder, potentially due to more evenly distributed compression in the spermatheca during eggshell formation (Fig. 8F, F’, H). Together, these data suggest that TXBP-3 and SPV-1 both contribute to the coordination of neck, bag, and valve contractility for maintaining embryo shape and preventing the embryos from severing during spermathecal transit.

### TXBP-3 and SPV-1 regulate actin fiber bundle alignment

To further investigate the cellular mechanism behind the transit defects, we observed the organization of the actin cytoskeleton (Fig. 9) using animals expressing F-tractin::GFP in the spermatheca. F-tractin is a peptide that binds actin (Melak et al., 2017). In WT backgrounds, F-tractin::GFP-expressing animals on both OP50 (Fig. 9A) and *empty*(RNAi) in HT115(DE3) (Fig. 9F) exhibit actin fibers organized into highly ordered, parallel bundles that are circumferentially aligned to facilitate coordinated contraction.

**Figure 9.**
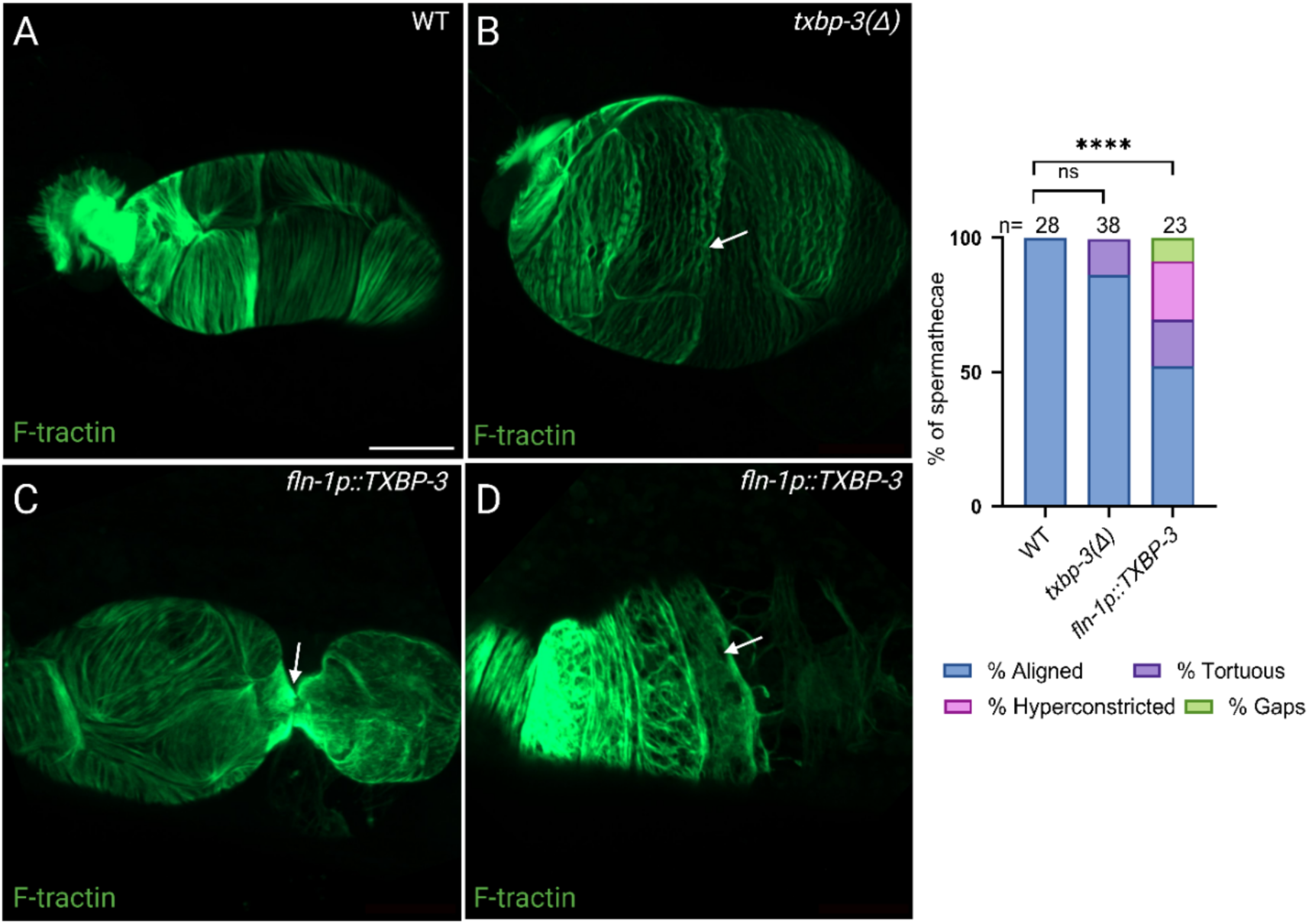
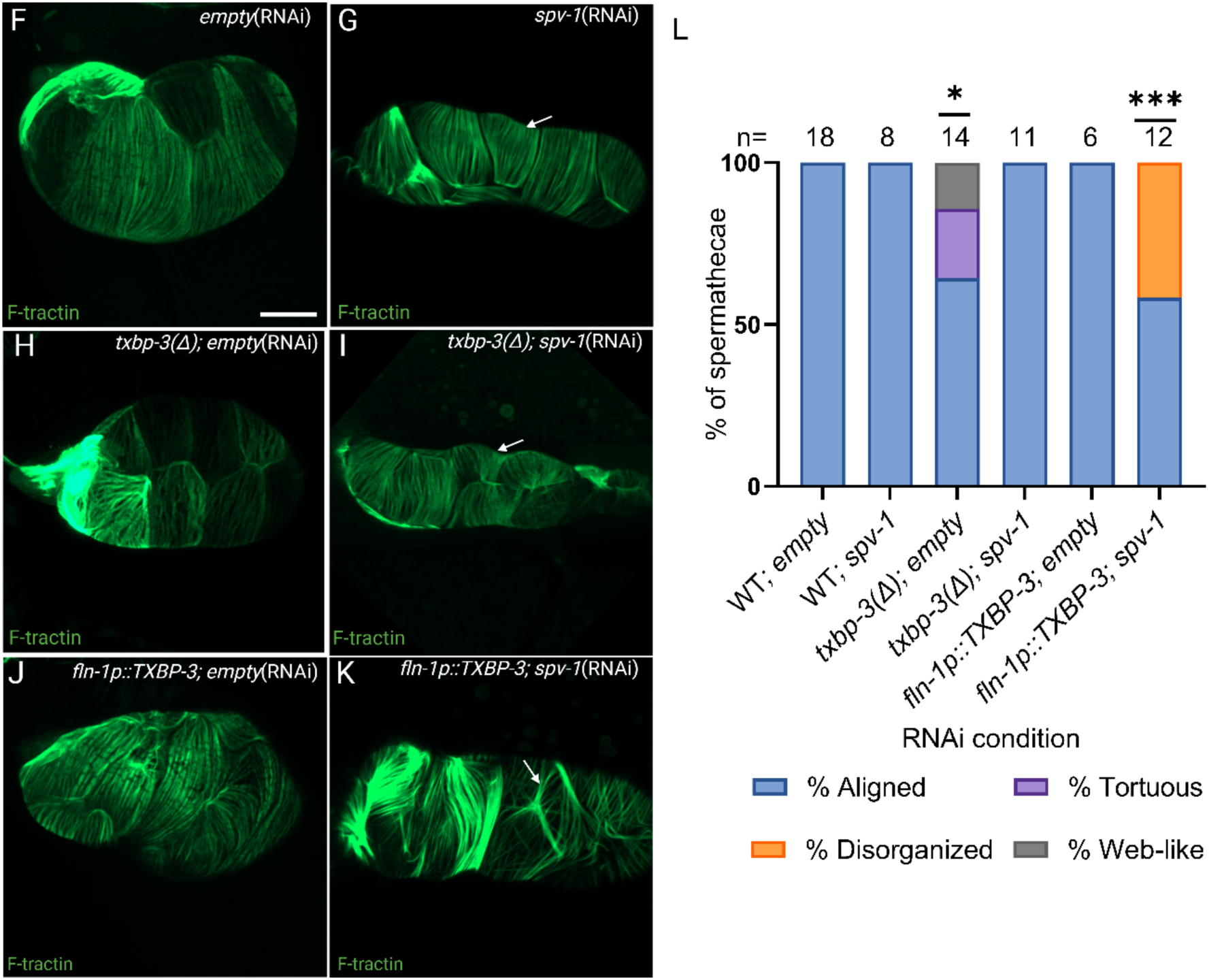
TXBP-3 and SPV-1 are essential for organized F-actin architecture in the spermatheca. (A-D) Representative confocal images of F-tractin::GFP in (A) WT, (B) *txbp-3(Δ)* (arrow: tortuosity), and (C-D) *fln-1p::*TXBP-3 (arrows: hyperconstricted bag and actin gaps). (E) Stacked bar chart of actin phenotypes showing aligned (blue), tortuous (purple), hyperconstricted (pink), and gaps (green) in actin across genotypes. (F-K) Representative F-tractin::GFP images across indicated RNAi conditions: (F) WT; *empty*(RNAi), (G) WT; *spv-1*(RNAi) (arrow: narrow spermatheca), (H) *txbp-3(Δ)*; *empty*(RNAi), (I) *txbp-3(Δ)*; *spv-1*(RNAi) (arrow: narrow spermatheca), (J) *fln-1p::*TXBP-3; *empty*(RNAi), and (K) *fln-1p::*TXBP-3; *spv-1*(RNAi) (arrow: clumped actin bundles). (L) Stacked bar chart of actin fiber phenotypes aligned (blue), tortuous (purple), disorganized (orange), and web-like (gray) phenotypes across the indicated genotypes. Scale bar = 20 µm. Statistical significance for categorical frequencies was determined using Fisher’s exact test (\**p* < 0.05, \*\*\**p* < 0.001, \*\*\*\**p* < 0.0001).

In contrast, disruptions to TXBP-3 resulted in specific cytoskeletal abnormalities. Loss of *txbp-3* resulted in tortuous (wavy) actin fibers in some spermathecae (13.2%, 5/38, ns vs. WT; Fig. 9B, E). Overexpression of *fln-1p::*TXBP-3 led to greater disorganization, including tortuosity (17.4%, 4/23), hyperconstriction (21.7%, 5/23; Fig. 9C), and gaps between fiber bundles (8.7%, 2/23; Fig. 9D, E; Table S3).

Through inhibition of RHO-1, SPV-1 regulates actomyosin tension. *spv-1*(RNAi) is known to increase tension on basal fibers, although it does not significantly disrupt the parallel fiber bundle alignment (Tan & Zaidel-Bar, 2015). Consistent with these findings, we observed that in *spv-1*(RNAi) animals, fiber bundles maintained their alignment (Fig. 9G). While most *txbp-3(Δ); empty*(RNAi) single mutants maintained aligned fibers (64.3%, 9/14; Fig. 9H), some showed tortuous fibers (21.4%, 3/14) or a disorganized, web-like appearance (14.3%, 2/14). The *txbp-3(Δ)* animals treated with *spv-1*(RNAi) were phenotypically similar to WT animals treated with *spv-1*(RNAi), with parallel actin fiber bundle alignment (100%, 11/11) (Fig. 9I; Table S3).

The *fln-1p::TXBP-3* animals exhibited aligned actin fibers on *empty*(RNAi) (Fig. 9J); however, combining *spv-1* depletion with TXBP-3 overexpression led to a loss of alignment and clumping into thicker bundles (41.7%, 5/12; Fig. 9K, L). These thicker bundles may exert increased tension, damaging oocytes during transit. These findings demonstrate that coordinated SPV-1 and TXBP-3 activity is essential for the actin structural integrity required for ovulation. Taken together, our results suggest TXBP-3 acts to activate UNC-43 and, in parallel with the Rho signaling pathway, to promote spermathecal and sp-ut valve contractility (Fig. 10).

**Figure 10.**
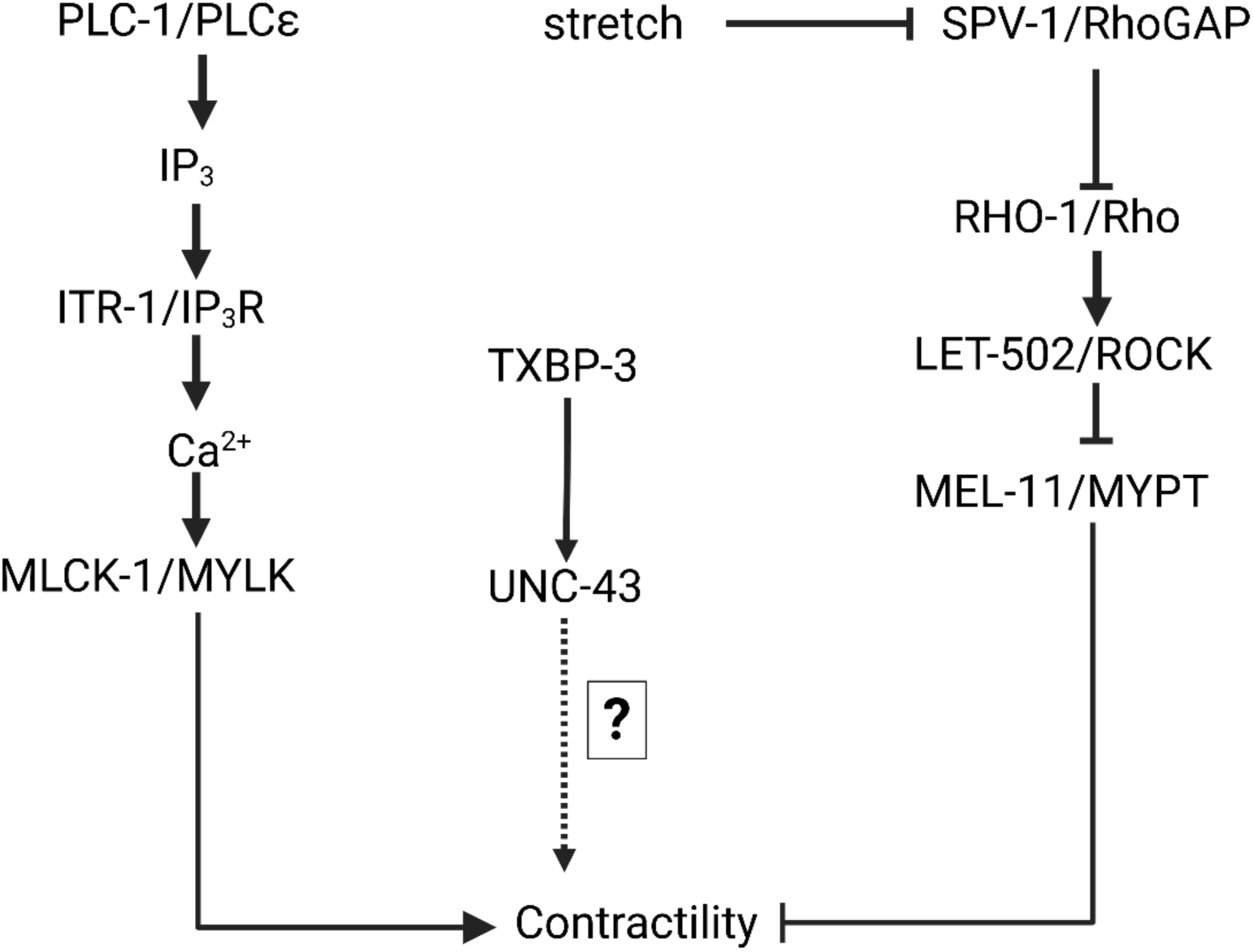
A working model of the spermathecal contractility network. We propose that TXBP-3 works in parallel with the Rho pathway to drive tissue contraction. TXBP-3 activates UNC-43. Both the calcium pathway and the Rho pathway converge to control spermathecal contractility.

## Discussion

In the *C. elegans* reproductive system, the precise coordination of contractile forces is required for successful oocyte transit. We identified TXBP-3, a highly conserved single-PDZ domain protein expressed in contractile tissues, as a regulator of spermathecal contractility. While baseline transit appears normal in *txbp-3(knu748)* mutants, sensitizing the tissue with the fluorescent calcium sensor GCaMP3 unmasks hidden defects, including delayed distal neck closure and premature opening of the sp-ut valve. Conversely, overexpressing TXBP-3 (*fln-1p::*TXBP-3) drives a hypercontractile state characterized by oocyte severing and embryo trapping. These failures suggest that successful ovulation relies on the precise coordination of the neck, bag, and valve.

TXBP-3 may work at least partially through activating UNC-43/CaMKII to regulate spermathecal contractility. Binding interactions between the C-terminus of UNC-43 and the TXBP-3 PDZ domain could induce a conformational change that promotes the open, active state of the kinase. Supporting this, levels of active, autophosphorylated CaMKII (Thr286) were significantly reduced in the *txbp-3* deletion mutant. Upon calcium influx, Ca^2+^/CaM binding triggers autophosphorylation at Thr286 (Miller & Kennedy, 1986), inducing calmodulin trapping to sustain kinase activity (Meyer et al., 1992). TXBP-3 binding may enhance this process, where initial activation of UNC-43 allows adjacent CaMKII multimer subunits to activate one another continuously (Hanson et al., 1994; Chao et al., 2011).

While the downstream targets of UNC-43 in the spermatheca are unknown, mammalian CaMKII interacts with several proteins. In cardiac muscle, CaMKII phosphorylates the ryanodine receptor RyR2 to enhance sarcoplasmic reticulum Ca^2+^ release (Maier et al., 2003). If UNC-43 acts similarly on the *C. elegans* ryanodine receptor ortholog UNC-68, this could help explain the excess Ca^2+^ release that accompanies sp-ut valve hypercontractility during TXBP-3 overexpression. Additionally, CaMKII binds α-actinin to anchor the kinase to F-actin and regulate its activity (Walikonis et al., 2001; Jalan-Sakrikar et al., 2012). If UNC-43 anchors ATN-1/α-actinin, disrupting this tether might help explain the actin disorganization seen when TXBP-3 levels are altered.

Spermathecal contractility is governed by two parallel pathways, a Ca^2+^-dependent pathway and a Rho-dependent pathway that converge to phosphorylate and activate non-muscle myosin II (Sethi et al., 2017; Bouffard et al., 2019). Loss of TXBP-3 results in lower and more variable Ca^2+^ signal, likely contributing to reduced contractility. Conversely, TXBP-3 overexpression can lead to failure of embryo exit, accompanied by strong Ca^2+^ pulses in the bag and valve, which appear to result in a tightly contracted sp-ut valve. Depletion of UNC-43 can also lead to embryo trapping and strong Ca^2+^ pulses. We have observed similar strong Ca^2+^ pulses upon embryo transit failures in a variety of different contexts, including depletion of the actin crosslinker FLN-1/filamin (Kovacevic et al., 2013), the Delta/Serrate/LAG-2 (DSL) family ligand APX-1 (McGovern et al., 2018), the transcription factor ATF-6 (Burkewitz et al., 2020), and the ZP-like protein PES-8 (Sadeghian & Cram, 2026). This suggests these ‘failure to exit’ Ca^2+^ pulses might be a response to the trapped embryo rather than an indication that the protein in question directly regulates Ca^2+^ release.

Tension in the spermatheca is controlled by the opposing actions of RhoGEFs, which activate RHO-1 to promote contraction, and RhoGAPs, which deactivate RHO-1 to permit relaxation (Tan & Zaidel-Bar, 2015; Avivi Kela et al., 2023). Depletion of the RhoGAP SPV-1 triggers RHO-1-driven hypercontractility in the distal spermatheca, rapidly pushing the oocyte through a tight sp-ut valve and causing high rates of pinching and oocyte fragmentation (Tan & Zaidel-Bar, 2015). In contrast, TXBP-3 overexpression drives hypercontractility at both the entrance neck and the exit valve. The entrance neck squeezes shut prematurely, severing the oocyte, and the exit valve locks as a closed sphincter. Removing pro-contractile TXBP-3 partially relieves the excess Rho-driven tension caused by loss of SPV-1. Conversely, activating both hypercontractile forces simultaneously by overexpressing TXBP-3 in the *spv-1(Δ)* mutant results in a mechanical failure. Early neck closure traps or severs the oocyte at the entrance, which precludes observation of potential exit-related sp-ut valve defects, decreasing overall fragmentation without resulting in a true genetic rescue.

These interactions are also reflected in the actin cytoskeleton. Successful ovulation requires an aligned, parallel actin bundle architecture in the spermatheca. Both loss and overexpression of TXBP-3 disrupt this organization, producing fiber tortuosity, web-like disorganization, hyperconstriction, and gaps. At the level of cytoskeletal organization, TXBP-3 again acts as a counterbalance to SPV-1: depleting SPV-1 in *txbp-3(Δ)* mutants eliminates the web-like actin disorganization, but causes the tissue to become tightly constricted, similar to *spv-1* mutants alone, whereas combining TXBP-3 overexpression with *spv-1*(RNAi) leads to thicker and potentially hypercontractile actin bundles.

In this study, we observed that HT115(DE3) RNAi food affects *txbp-3(Δ)* and TXBP-3 overexpression-related phenotypes, and therefore, we compared effects only between animals reared on the same diet. *C. elegans* is generally reared on the *E. coli* strain OP50 (Brenner, 1974). OP50 and HT115(DE3) differ in their nutritional content and have variable effects on nematode physiology, survival, and reproduction (Reinke et al., 2010). These dietary differences have been linked to factors such as vitamin B12 deficiency, compromised mitochondrial homeostasis, and redox stresses (Pang & Curran, 2014; Revtovich et al., 2019; Nair et al., 2022). The mechanism underlying the effect of food source on spermathecal contractility and calcium signaling coordination is a rich topic for future investigation.

In summary, this study adds to our knowledge of how PDZ domain proteins regulate contractility *in vivo*, and illustrates the importance of coordinated calcium dynamics, regulation and alignment of contractile actomyosin bundles, as well as cooperation between cells in contractile tissues. Given the strong conservation of TXBP-3, homologous proteins such as TAX1BP3 may play similar roles in other animals including humans.

## Materials and Methods

### Plasmid Construction

To examine *txbp-3* expression, we constructed both a transcriptional reporter pUN1500 (*txbp-3p::GFP*) and a translational fusion plasmid pUN846 (*txbp-3p::TXBP-3::GFP*), which were microinjected into N2 (Table 3). pUN846 was generated by amplifying the genomic region of *txbp-3* (C45G9.7) and the pPD95.77 backbone (Andrew Fire) using NEBuilder HiFi DNA Assembly. A two-step PCR strategy was used; *txbp-3* was first amplified with gene-specific primers, followed by a second PCR to generate NEBuilder-compatible overlaps. Gene-specific primers (Table 1) were used to amplify the *txbp-3* genomic region, and vector-specific primers (Table 1) were used to linearize the vector backbone. The resulting insert, comprising the *txbp-3* promoter and the full-length genomic coding sequence in-frame with GFP, was assembled into the pPD95.77 backbone to produce a C-terminal translational fusion.

**Table 1.**
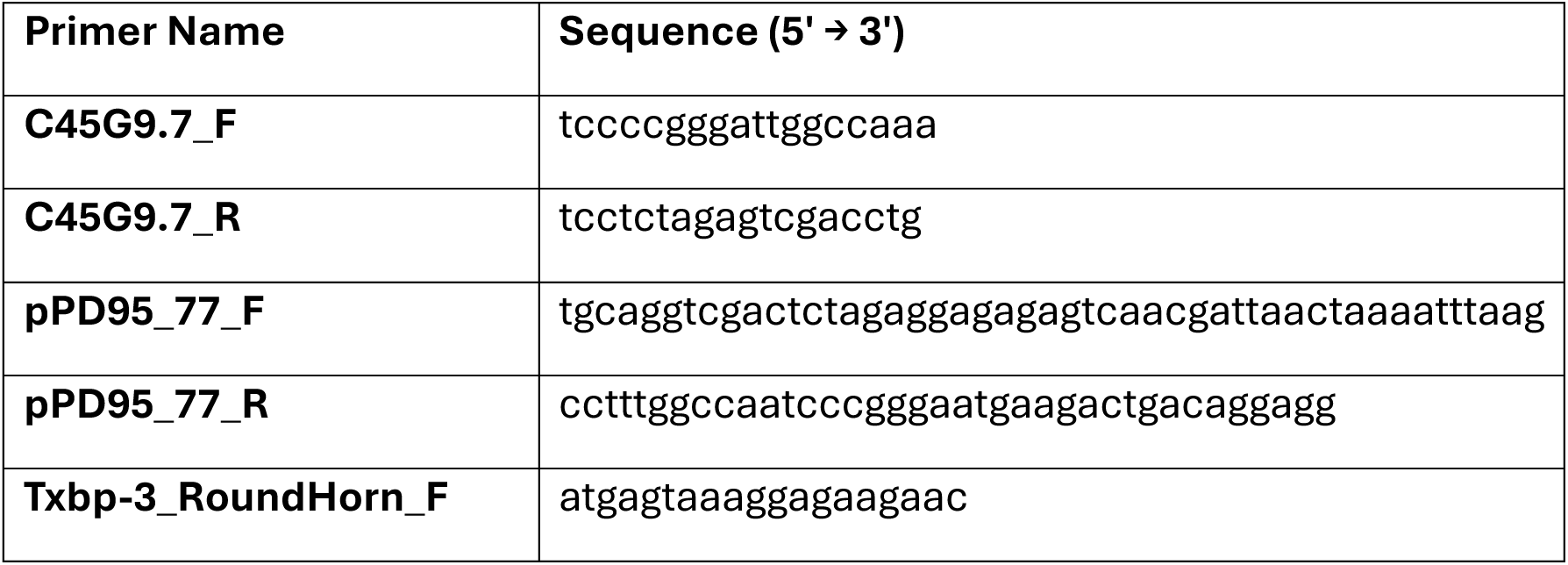
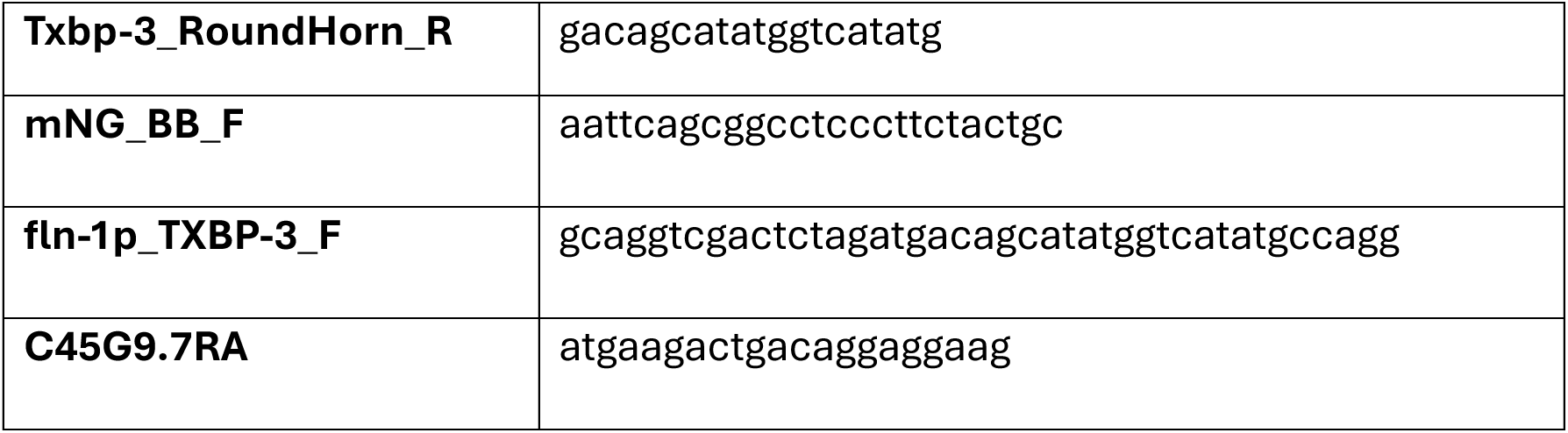
Primers for plasmid construction.

To generate the transcriptional reporter plasmid, pUN1500, we applied the Round-the-Horn PCR method to amplify the *txbp-3* promoter and GFP region from pUN846 while excluding the *txbp-3* coding sequence. The resulting amplicon was re-ligated to generate a promoter-only GFP fusion using specific primers for this deletion (Table 1).

To overexpress TXBP-3 in the spermatheca, we generated pUN1881 (*fln-1p::TXBP-3::fln-1 3′UTR*). The *txbp-3* genomic coding sequence was first inserted into pUN71 (*fln-1p::GFP*) to create the intermediate plasmid pUN1861 (*fln-1p::TXBP-3::GFP*). The GFP coding sequence was then removed by inverse PCR amplification of pUN1861 with primers mNG_BB_F and C45G9.7RA (Table 1), followed by T4 polynucleotide kinase (PNK) treatment and self-ligation. The construct was confirmed by sequencing and injected into N2 animals with *myo-2p::mCherry* as a co-injection marker to generate the extrachromosomal array *xbEx2518* (Table 3).

### *C. elegans* Strains and Culture

*Caenorhabditis elegans* strains were cultured on NGM plates consisting of 50 mM NaCl, 2.5 g/L Peptone, 17 g/L BD Bacto-Agar, 5 mL/L Nystatin, 1 mM CaCl₂, 1 mM MgSO₄, 0.5 mL/L of 10 mg/mL cholesterol, and 25 mM KPO₄, and fed with *E. coli* strains OP50 or HT115(DE3) at 23°C, following standard protocols (Brenner, 1974). The *C. elegans* strain COP1859 *txbp-3(knu748)* carries a 1005 bp deletion generated using CRISPR/Cas9 genome editing. This deletion removes the entire coding sequence of *txbp-3* along with a portion of its upstream promoter, resulting in a null allele. The strain was generated by Knudra Transgenics (now In Vivo Biosystems) and backcrossed four times to N2 to eliminate background mutations. Several lines were generated through standard genetic crosses. Genotypes of cross progeny were confirmed by PCR amplification of deletion alleles and visual screening for fluorescent markers. A complete list of strains used and generated in this study is provided in Table 2. Injection mix compositions for transgenic lines are listed in Table 3.

**Table 2.** Strain List.

| **Strain** | **Genotype / Description** | **Source** |
| --- | --- | --- |
| **N2** | WT (Bristol) | CGC |
| **COP1859** | *txbp-3(knu748)* | In Vivo Biosystems |
| **NM1378** | *unc-43(js125)* | CGC |
| **RB976** | *rhgf-1(ok880)* | CGC |
| **RB1353** | *spv-1(ok1498)* | CGC |
| **UN1417** | *xbIs1417[fln-1p::GCaMP3]* | Cram lab |
| **UN1896** | *xbEx1896 [txbp-3p::TXBP-3::GFP]* | Cram lab |
| **UN1948** | *txbp-3(knu748); xbIs1417 [fln-1p::GCaMP3]* | Cram lab |
| **UN2221** | *xbIs2221[fln-1p::F-tractin::GFP; rol-6]* | Cram lab |
| **UN2426** | *txbp-3(knu748); xbIs2221[fln-1p::F-tractin::GFP; rol-6]* | Cram lab |
| **UN2506** | *xbEx2506[txbp-3p::GFP]* | Cram lab |
| **UN2507** | *rhgf-1(ok880); txbp-3(knu748)* | Cram lab |
| **UN2518** | *xbEx2518[fln-1p::TXBP-3; myo-2p::mCherry]* | Cram lab |
| **UN2529** | *spv-1(ok1498); txbp-3(knu748)* | Cram lab |
| **UN2530** | *xbIs2221[fln-1p::F-tractin::GFP; rol-6]; xbEx2518[fln-1p::TXBP-3; myo-2p::mCherry]* | Cram lab |
| **UN2531** | *spv-1(ok1498);xbEx2518[fln-1p::TXBP-3; myo-2p::mCherry]* | Cram lab |
| **UN2541** | *xbIs1417[fln-1p::GCaMP3];xbEx2518[fln-1p::TXBP-3; myo-2p::mCherry]* | Cram lab |
| **UN2542** | *rhgf-1(ok880); xbEx2518[fln-1p::TXBP-3; myo-2p::mCherry]* | Cram lab |

**Table 3.** Generation of Transgenic Lines via Microinjection.

| **Strain** | **Host Strain** | **Plasmid Construct** | **Injection Mix Composition** |
| --- | --- | --- | --- |
| **UN1896** | N2 | pUN846 (*txbp-3p::TXBP-3::GFP*) | pUN846 (20 ng/μL); 1kb DNA Ladder (50 ng/μL) |
| **UN2506** | N2 | pUN1500 (*txbp-3p::GFP*) | pUN1500 (20 ng/μL); 1kb DNA Ladder (50 ng/μL) |
| **UN2518** | N2 | pUN1881 (*fln-1p::TXBP-3*) | pUN1881 (20 ng/μL); *myo-2p::mCherry* (2 ng/μL); 1kb DNA Ladder (50 ng/μL) |

### RNA Interference (RNAi)

RNAi was conducted via the feeding method, as previously described by Timmons & Fire (1998). *E. coli* HT115(DE3) carrying the dsRNA construct of interest was grown overnight in Luria Broth (LB) containing 40 µg/mL ampicillin. NGM plates supplemented with 25 µg/mL carbenicillin and 1 mM IPTG were seeded with 200 µL of bacterial culture and incubated at room temperature for 24-72 hours to induce dsRNA expression. The *empty* L4440 vector (pPD129.36) served as the negative control for all assays.

To synchronize the populations, embryos were isolated from gravid adults using alkaline hypochlorite treatment (Hope, 1999) and subsequently washed three times in M9 buffer (22 mM KH₂PO₄, 42 mM Na_2_HPO₄, 86 mM NaCl, and 1 mM MgSO₄). Synchronized embryos were transferred directly to RNAi plates and incubated at 23°C for 60-64 hours, allowing worms to reach the young adult stage before analysis.

### Immunofluorescence of Dissected Gonads

To visualize phospho-UNC-43 levels, approximately 100 young adult hermaphrodites per condition were picked into M9 buffer on a watch glass. Gonads were rapidly dissected and immediately fixed in a final concentration of 2% neutral-buffered paraformaldehyde for 2 hours at 4°C in a humidified chamber. Following fixation, the paraformaldehyde was carefully removed, and the tissues were treated with pre-chilled (-20°C) methanol for 5 minutes. Samples were then transferred to glass graduated tubes, and the methanol was replaced with three consecutive washes in PBST (1X PBS containing 0.1% Tween-20), allowing the tissues to settle by gravity between washes. To minimize non-specific binding, gonads were blocked in 1% Bovine Serum Albumin (BSA) in PBST for 1 hour.

Samples were then incubated overnight at 4°C with a rabbit monoclonal anti-phospho-CaMKII (Thr286) primary antibody (clone HL3570, GeneTex, catalog no. GTX641517) diluted 1:250 in the blocking solution. The following day, gonads were washed three times in PBST and incubated for 2 hours with a donkey anti-rabbit DyLight 488-conjugated secondary antibody (Invitrogen) diluted 1:500 in blocking solution. After two final washes in PBST, the samples were mounted on 3% agarose pads for visualization. Wide-field fluorescence and Differential Interference Contrast (DIC) imaging were conducted using a Nikon Eclipse microscope equipped with a 60x oil-immersion objective and a Spot RT39M5 camera. To ensure quantitative comparability, all imaging parameters and exposure times were standardized across all experimental strains and negative controls.

### Time Point Metrics

To synchronize young adult worms for transit assays, gravid adults were lysed with an alkaline hypochlorite solution to release eggs, which were washed three times with M9 buffer, then placed on seeded NGM plates and grown at 23°C for 52-54 hours to reach the young adult stage. Imaging was performed on a Nikon Eclipse microscope with a 60x oil-immersion lens using SPOT Advanced software (Version 5.3.5) and a charge-coupled device camera.

To characterize ovulation dynamics and embryo transit through the spermatheca, young adult hermaphrodites were immobilized on 3% agarose pads prepared with a 1:2 dilution of M9 buffer supplemented with 0.05 µm Polybead microspheres. Time-lapse imaging was performed using DIC microscopy at 1 frame per second (1 Hz). For each ovulation event, four key time points were recorded: (1) oocyte entry onset, defined as the moment the spermatheca begins to move over the incoming oocyte; (2) distal spermathecal closure, when the oocyte is fully enclosed within the spermatheca; (3) spermathecal-uterine (sp-ut) valve opening, when embryo exit begins; and (4) sp-ut valve closure, when the embryo is fully expelled into the uterus and the valve re-constricts.

These time points were used to calculate three kinetic metrics: entry time (from the onset of entry to full enclosure), dwell time (the duration of complete embryo enclosure), and exit time (from the opening of the sp-ut valve to complete expulsion). All image sequences were analyzed using ImageJ (Fiji).

### Embryo Shape Assay

To quantify embryo morphology, five gravid adult hermaphrodites maintained at 23°C were transferred to a 3% agarose pad equilibrated with 1 µL of sodium hypochlorite solution (5.65-6% active chlorine; Fisher Chemical). After a 2-minute incubation to dissolve the cuticle and facilitate embryo release from the uterus, samples were rinsed with 2 µL of M9 buffer and secured with a coverslip. To prevent morphological changes during imaging, all images were acquired within 5 minutes of coverslip placement under non-fixative conditions.

Imaging was performed using Differential Interference Contrast (DIC) microscopy equipped with a 60x oil-immersion objective. Analysis was restricted to multicellular embryos captured across multiple fields of view; any embryos that were tilted, overlapping, or partially obscured were excluded. Embryo boundaries were manually segmented in ImageJ (Fiji) using the freehand selection tool. The following morphological parameters were extracted for each embryo: area (µm²), roundness (4 × area / (π × major axis²)), and aspect ratio (major axis/minor axis). A minimum of 68 embryos were analyzed per condition across three independent biological replicates, each processed on a separate day.

### Wide-field Fluorescence and Live Calcium Imaging

Differential Interference Contrast (DIC) and wide-field fluorescence imaging were conducted using a Nikon Eclipse microscope equipped with a 60x oil-immersion objective and a Spot RT39M5 camera (Diagnostic Instruments, Sterling Heights, MI, USA). Fluorescence was excited with a Nikon Intensilight C-HGFI 130 W mercury lamp, controlled by a SmartShutter (Sutter Instruments, Novato, CA, USA). For imaging, young adult animals were directly transferred from plates onto slides. To capture still images, animals were immobilized on 3% agarose pads with 0.1 M sodium azide. For time-lapse movies, animals were immobilized with 0.05 µm polystyrene polybeads (Polysciences Inc., Warrington, PA, USA) diluted 1:2 in M9 buffer.

Ca^2+^ dynamics were measured using the GCaMP3 sensor. All time-lapse images were captured at 1 frame per second (1 Hz). Movies obtained with the Spot RT39M5 camera used an exposure time of 20 ms and a gain of 8 (captured at 2448×2048 pixels). Data collection was restricted to the first three ovulations per animal, with a strong preference for the first ovulation. All imaging parameters were standardized across experimental conditions, and files were saved as 8-bit Tagged Image File Format (TIFF) files.

All image processing was conducted using custom macros in ImageJ (Kovacevic et al., 2013). Images were oriented with the sp-ut valve on the right side of the frame and registered to reduce body movement artifacts. A 942 × 471-pixel ROI encompassing the entire spermatheca was used to measure the GCaMP3 signal. Ca^2+^ pixel intensity (F) was normalized to the average pixel intensity of the first 30 frames before the start of ovulation (F₀) and plotted against time.

Data analysis and graphing were performed using MATLAB and GraphPad Prism. Local maxima in the time series with a minimum prominence of 0.1 (F/F₀) and minimum width of 5 s were then identified using the MATLAB ‘findpeaks’ command. These standardized settings were used to analyze all Ca^2+^ traces. To determine the rate at which the Ca^2+^ signal changed with time, kymograms were generated using an ImageJ macro that calculated the average pixel intensity of each column of a frame and condensed it down to one line per frame of the time-lapse image. Every frame of the time-lapse image was stacked to visualize Ca^2+^ dynamics of representative ovulations in both space and time. The Fire lookup table color scale was applied to the kymograms using ImageJ. All MATLAB and ImageJ scripts are available upon request.

### Confocal Imaging

For fixed sample imaging, synchronized animals were grown at 23°C for 60-64 hours. They were then fixed in 4% formaldehyde (in 1X PBS) for 20 minutes at room temperature, washed three times in 1X PBS, and mounted on 3% agarose pads.

All imaging was performed on a Zeiss LSM 800 confocal microscope equipped with a Plan-Apochromat 63x/1.40 NA oil DIC M27 objective. The setup was driven by ZEN software. GFP was excited at 488 nm. For 3D reconstructions, z-stacks were acquired at 0.33 µm intervals, with 40 slices per volume. Orthogonal projections were generated in ZEN.

### Image Analysis and Statistics

All image processing and quantitative analysis were done using Fiji (ImageJ 2.3.0/1.53f). Statistical analysis was performed with GraphPad Prism 9.2.0. For continuous measurements (e.g., transit times, Ca^2+^ peak parameters, and embryo area, aspect ratio, and roundness), genotypes were compared by ordinary one-way ANOVA followed by Šídák’s test for comparisons among selected pairs of genotypes, Dunnett’s test for comparisons of each genotype to a single control, or Tukey’s test for comparisons among all genotypes; the test used for each dataset is indicated in the corresponding figure legend. Fisher’s exact test was used for categorical outcomes. Unless noted, data are shown as mean ± SD. Significance is denoted: **** *p* < 0.0001; *** *p* < 0.001; \*\**p* < 0.01; * *p* < 0.05; ns if *p* ≥ 0.05.

## Acknowledgements

We thank Avery Lord for beginning this project and members of the Cram and Apfeld labs for helpful discussions. Some strains were provided by the CGC, which is funded by NIH Office of Research Infrastructure Programs (P40 OD010440). This work was supported by grants from the National Institutes of Health National Institute of General Medical Sciences (GM110268) and the National Science Foundation – Binational Science Foundation (2419801) to E.J.C.

## Abbreviations

CaMKII: Calcium/Calmodulin-dependent Protein Kinase II
CGC: *Caenorhabditis* Genetics Center
DAG: Diacylglycerol
DIC: Differential Interference Contrast
GAPs: GTPase-activating proteins
GCaMP3: GFP-calmodulin (CaM) and M13 Peptide Version 3
GEFs: Guanine nucleotide exchange factors
IP3: Inositol trisphosphate
MFI: Mean fluorescence intensity
MLCK-1: Myosin light chain kinase
NGM: Nematode growth medium
PDZ: PSD-95, Dlg, and ZO-1 domain
pLDDT: Predicted local distance difference test
RNAi: RNA interference
RhoGAP: Rho GTPase-Activating Protein
RhoGEF: Rho Guanine Nucleotide Exchange Factor
RyR2: Ryanodine receptor 2
sp-ut: Spermathecal-uterine valve
TAX1BP3: Tax1 Binding Protein 3 (mammalian ortholog)
TEAD1: Transcriptional Enhanced Associate Domain 1
TIP-1: Tax-interacting Protein 1 (mammalian ortholog)
TRPV4: Transient Receptor Potential Cation Channel Subfamily V Member 4
TXBP-3: Tax1 Binding Protein 3
YAP: Yes-Associated Protein

