## Supplemental Tables and Figures for "The single-PDZ domain protein TXBP-3 regulates contractility in the *C. elegans* spermatheca"

**Supplementary Movies:**

**Movie S1: WT oocyte transit.** Time-lapse DIC imaging of a normal oocyte transit in a WT animal fed on OP50. The oocyte enters from the right and undergoes a successful, coordinated entry, dwell, and exit through the sp-ut valve.

**Movie S2: Oocyte transit in *txbp-3(Δ)* mutants.** Time-lapse DIC imaging of an oocyte transit in a *txbp-3(Δ)* mutant fed on OP50. The oocyte enters from the right and transit proceeds successfully, with entry, dwell, and exit dynamics that are comparable to WT animals.

**Movie S3: Oocyte transit in TXBP-3 overexpression animals.** Time-lapse DIC imaging of an oocyte transit in a *fln-1p::*TXBP-3 animal fed on OP50. The oocyte enters from the right. Because of TXBP-3-driven hypercontractility, the spermathecal neck closes prematurely, severing the oocyte into two pieces and leaving a fragment behind in the gonad arm.

**Movie S4: $Ca^{2+}$ dynamics during oocyte transit in a WT animal.** Time-lapse fluorescence imaging of GCaMP signal during a successful oocyte transit in a WT (*fln-1p::GCaMP*) animal. Oocyte entry triggers a rhythmic calcium wave that propagates from the distal neck toward the sp-ut valve.

**Movie S5: $Ca^{2+}$ dynamics during oocyte transit in a *txbp-3(Δ)* mutant.** Time-lapse fluorescence imaging of GCaMP signal during an oocyte transit in a *txbp-3(Δ)* mutant. A rhythmic calcium wave is visible, but the overall calcium signal intensity is weaker.

**Movie S6: $Ca^{2+}$ dynamics during oocyte transit in a TXBP-3 overexpression animal.** Time-lapse fluorescence imaging of GCaMP signal during an oocyte transit in a *fln-1p::*TXBP-3 overexpression animal. The oocyte fails to exit and becomes trapped, accompanied by a hyperactive calcium response with continuous, elevated calcium pulses within the spermatheca.

**Movie S7: Oocyte transit in a WT animal treated with *empty*(RNAi).** Time-lapse imaging of spermathecal $Ca^{2+}$ dynamics during a successful oocyte transit in a WT (*fln-1p::GCaMP*) animal fed on *empty*(RNAi) control.

**Movie S8: Oocyte transit in a WT animal treated with *unc-43*(RNAi).** Time-lapse imaging of spermathecal $Ca^{2+}$ dynamics during an abnormal oocyte transit in a WT (*fln-1p::GCaMP*) animal fed on *unc-43*(RNAi). The oocyte exhibits abnormal back-and-forth movement.

**Movie S9: Oocyte transit in *spv-1(Δ)* mutants.** Time-lapse DIC imaging of a successful oocyte transit in an *spv-1(Δ)* mutant fed on OP50. The oocyte enters from the right. While a subpopulation of these mutants exhibits valve tightness and oocyte severing, this representative video shows complete transit characterized by hypercontractility and rapid exit dynamics.

**Movie S10: Oocyte transit in *txbp-3(Δ); spv-1(Δ)* double mutants.** Time-lapse DIC imaging of a successful oocyte transit in a *txbp-3(Δ); spv-1(Δ)* double mutant fed on OP50. The oocyte enters from the right and successfully exits the spermatheca.

**Movie S11: Oocyte transit in *fln-1p::*TXBP-3*; spv-1(Δ)* animals.** Time-lapse DIC imaging of oocyte transit in a *fln-1p::*TXBP-3*; spv-1(Δ)* animal. The oocyte enters from the right. Hypercontractility at both the spermathecal neck and the sp-ut valve disrupts transit, resulting in mechanical defects such as the oocyte being severed.

#### Supplementary data

##### Table S1: Summary of oocyte transit outcomes and failure modes

| Genotype | n | Successful Transit | Neck Tight | Valve Tight | Oocyte Rupture | Trapped | Oocyte Pushback |
|---|---|---|---|---|---|---|---|
| N2 (WT) | 11 | 100% (11) | - | - | - | - | - |
| *txbp-3(Δ)* | 10 | 90.0% (9) | 10% (1) | - | - | - | - |
| *fln-1p::*TXBP-3 | 13 | 61.5% (8) | 23.1% (3) | 15.4% (2) | 15.4% (2) | 7.7% (1) | - |
| *fln-1p::GCaMP* (WT) | 11 | 100% (11) | - | - | - | - | - |
| *txbp-3(Δ); GCaMP* | 7 | 100% (7) | - | - | - | - | - |
| *fln-1p::*TXBP-3*; GCaMP* | 8 | 50% (4) | - | - | - | 25% (2) | 25% (2) |
| *rhgf-1(Δ)* | 8 | 87.5% (7) | 12.5% (1) | - | - | - | - |
| *txbp-3(Δ); rhgf-1(Δ)* | 9 | 100% (9) | - | - | - | - | - |
| *fln-1p::TXBP-3 ; rhgf-1(Δ)* | 10 | 80% (8) | 10% (1) | - | - | 10% (1) | - |
| *spv-1(Δ)* | 16 | 62.5% (10) | 6.3% (1) | 25% (4) | 6.3% (1) | - | - |
| *txbp-3(Δ); spv-1(Δ)* | 20 | 55% (11) | 35% (7) | 5% (1) | 10% (2) | 5% (1) | - |
| *fln-1p::TXBP-3; spv-1(Δ)* | 22 | 31.8% (7) | 45.5% (10) | 13.6% (3) | 4.5% (1) | 13.6% (3) | 9.1% (2) |

Categories are not mutually exclusive; some transits displayed more than one failure mode.

**Table S2: Quantification of embryo morphology**

| Genotype | Total (n) | Normal % (n) | Fragmented % (n) |
|---|---|---|---|
| N2 (WT) | 68 | 100% (68) | - |
| *spv-1(Δ)* | 76 | 75% (57) | 25% (19) **** |
| *txbp-3(Δ)* | 75 | 100% (75) | - |
| *txbp-3(Δ); spv-1(Δ)* | 75 | 86.7% (65) | 13.3% (10) ** |
| *fln-1p::*TXBP-3 | 75 | 80% (60) | 20% (15) **** |
| *fln-1p::*TXBP-*3; spv-1(Δ)* | 76 | 89.5% (68) | 10.5% (8) ** |

Statistical significance compared to WT was determined by Fisher's exact test (***p* < 0.01, *****p* < 0.0001).

**Table S3: Quantitative distribution of spermathecal F-actin phenotypes**

| Genotype | Total (n) | Aligned %(n) | Dis-organized %(n) | Tortuous %(n) | Hyperconstricted %(n) | Gaps %(n) | Web-like %(n) |
|---|---|---|---|---|---|---|---|
| *F-tractin* (WT) | 28 | 100% (28) | - | - | - | - | - |
| *F-tractin*; *txbp-3(Δ)* | 38 | 86.8% (33) | - | 13.2% (5) | - | - | - |
| *F-tractin; fln-1p::TXBP-3* | 23 | 52.2% (12) | - | 17.4% (4) | 21.7% (5) | 8.7% (2) | - |
| *F-tractin*; *empty*(RNAi) | 18 | 100% (18) | - | - | - | - | - |
| *txbp-3(Δ); empty*(RNAi) | 14 | 64.3% (9) | - | 21.4% (3) | - | - | 14.3% (2) |
| *fln-1p::TXBP-3; empty*(RNAi) | 6 | 100% (6) | - | - | - | - | - |
| *F-tractin*; *spv-1*(RNAi) | 8 | 100% (8) | - | - | - | - | - |
| *txbp-3(Δ); spv-1*(RNAi) | 11 | 100% (11) | - | - | - | - | - |
| *fln-1p::TXBP-3; spv-1*(RNAi) | 12 | 58.3% (7) | 41.7% (5) | - | - | - | - |

#### Supplementary figures

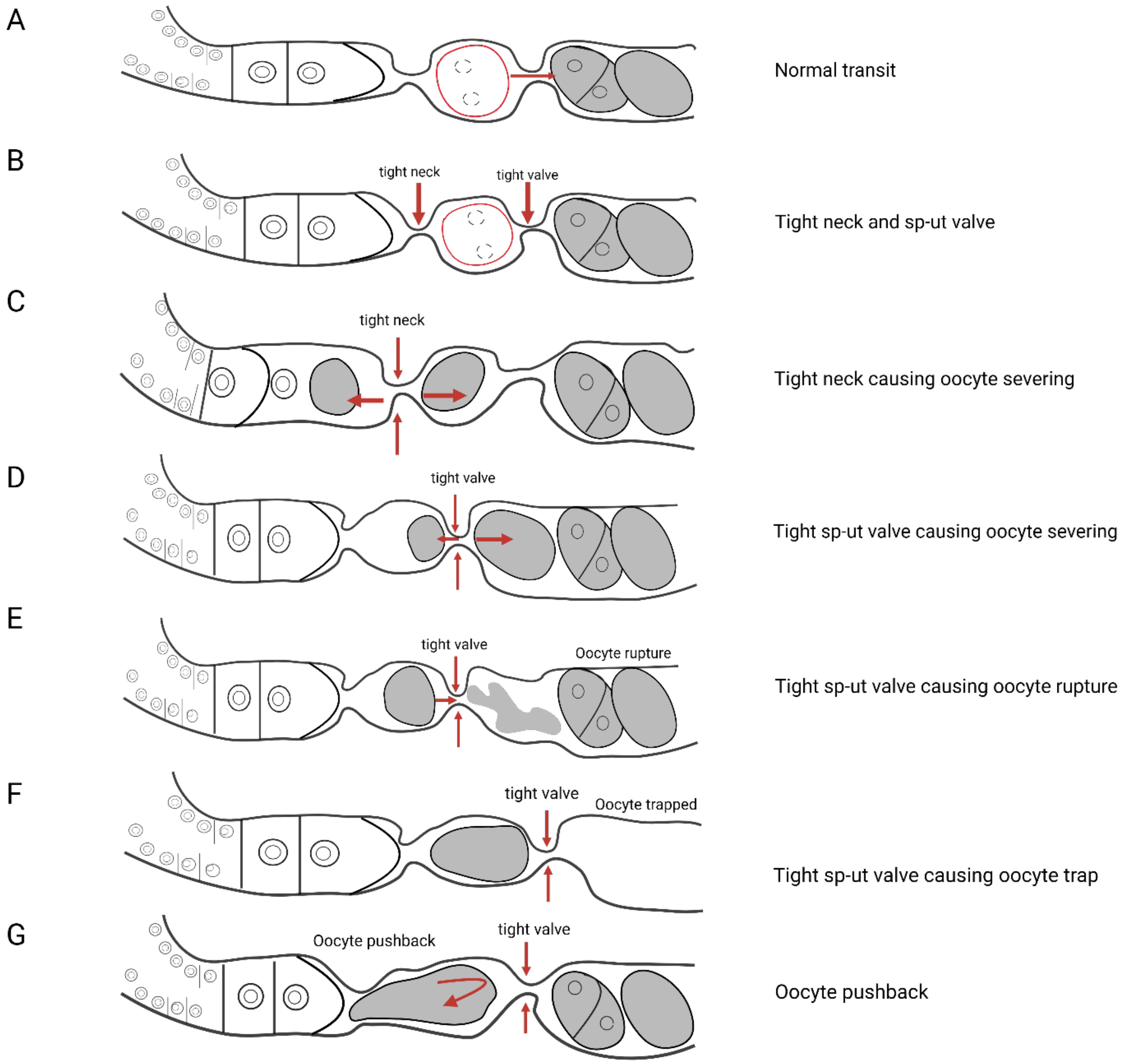


**Figure S1. Coordination of neck and sp-ut valve contractility is essential for spermathecal transit.** (A-G) Schematic illustrating normal oocyte transit and different mechanical failures observed during hypercontractility. (A) In a successful transit, the oocyte passes smoothly through the spermathecal neck and sp-ut valve. (B) Premature tightening of the neck and sp-ut valve. (C) Oocyte severing by the tight neck. (D) Oocyte severing by the tight sp-ut valve. (E) Oocyte rupture. (F) Oocyte trapped in the spermatheca. (G) Oocyte pushed back into the gonad arm.

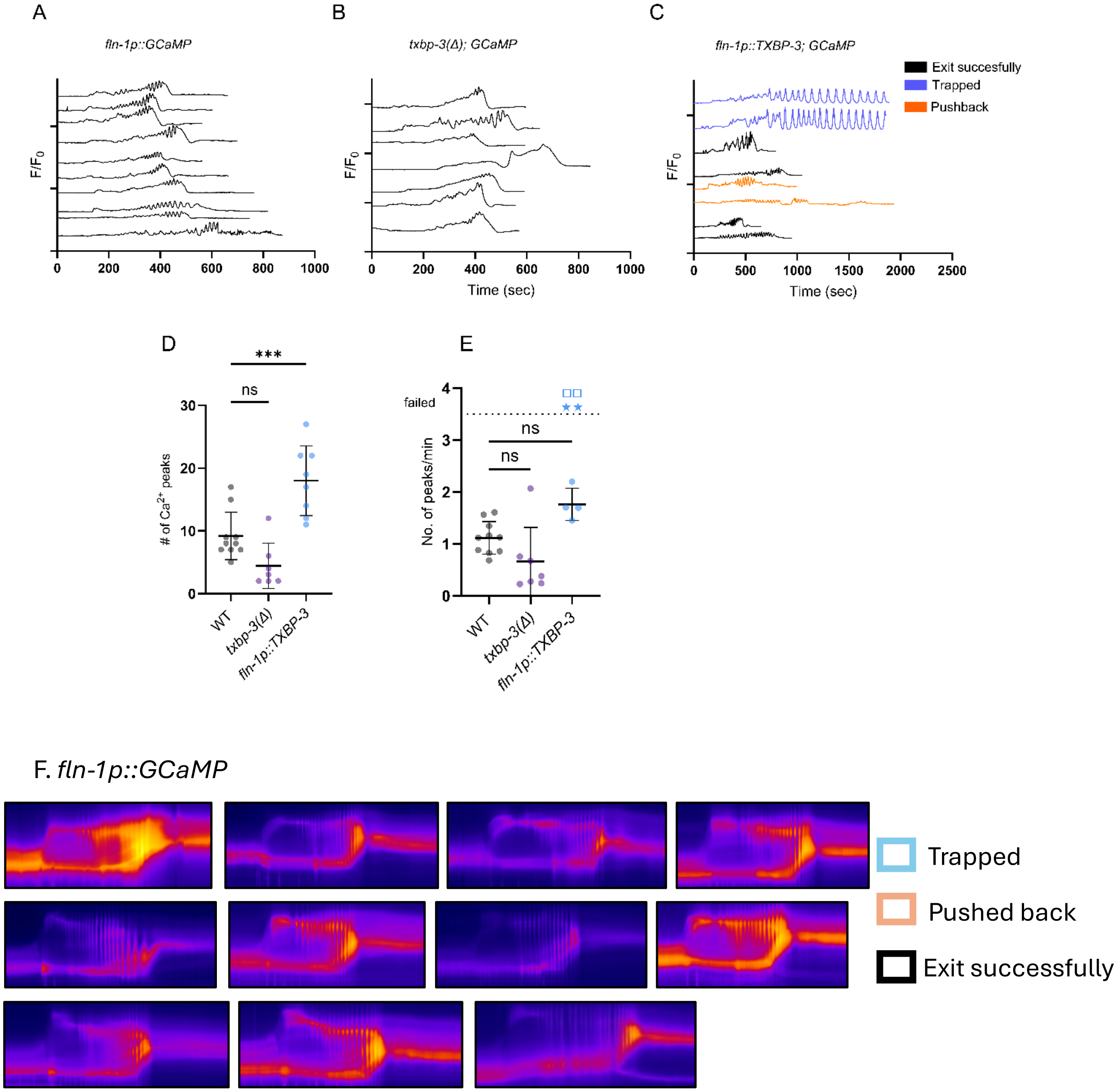
A
fln-1p::GCaMP
F/F0
0 200 400 600 800 1000
B
txbp-3(Δ); GCaMP
F/F0
0 200 400 600 800 1000
Time (sec)
C
fln-1p::TXBP-3; GCaMP
Exit succesfully
Trapped
Pushback
F/F0
0 500 1000 1500 2000 2500
Time (sec)
D
***
ns
30
20
10
0
### of Ca2+ peaks
WT
txbp-3(Δ)
fln-1p::TXBP-3
E
failed
4
3
2
1
0
ns
ns
No. of peaks/min
WT
txbp-3(Δ)
fln-1p::TXBP-3
F. fln-1p::GCaMP
Trapped
Pushed back
Exit successfully

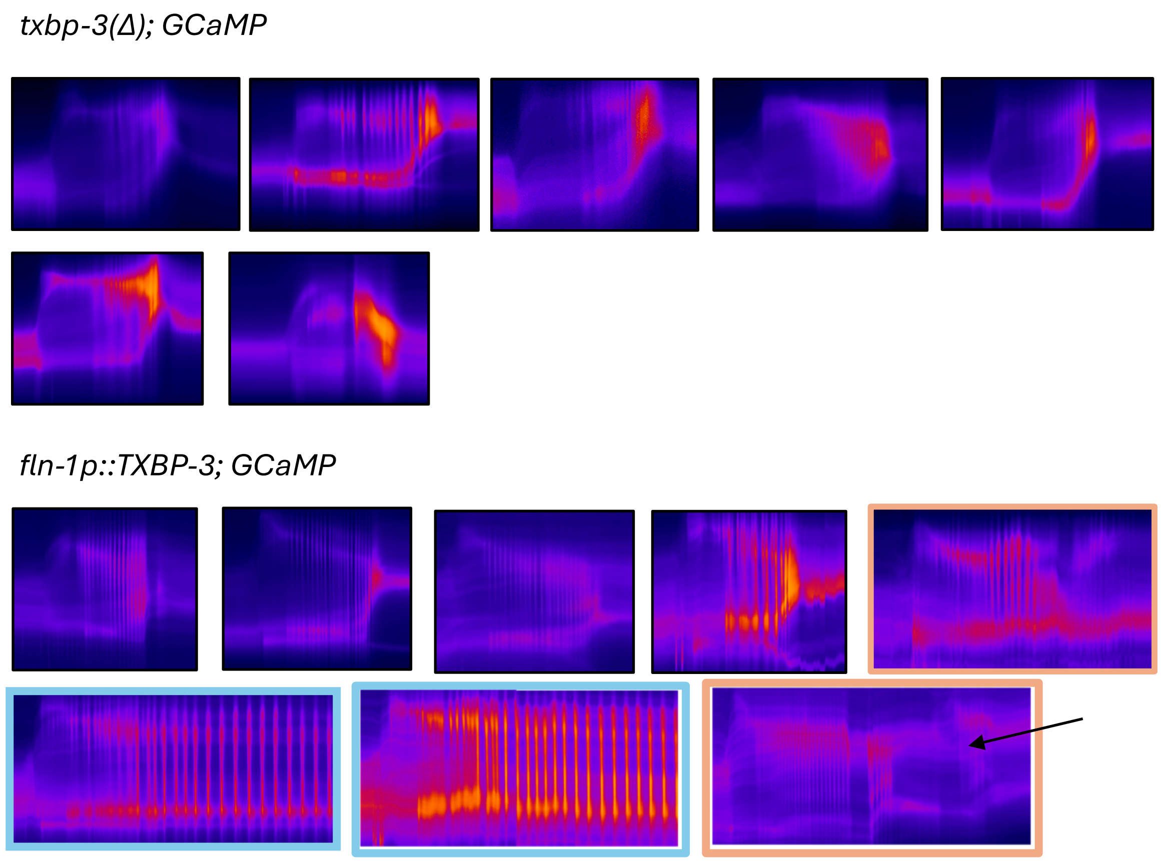


**Figure S2. 1D traces and kymograms of spermathecal calcium transients.** (A-C) Multiple representative 1D traces showing normalized $Ca^{2+}$ signal intensity ($F/F_0$) over time (sec) for individual oocyte transit events in (A) *fln-1p::GCaMP* (WT), (B) *txbp-3(Δ); GCaMP*, and (C) *fln-1p::*TXBP-3*; GCaMP* animals. In (C), individual traces are color-coded by transit outcome: successful exit (black), trapped oocyte (blue), and pushback (orange). Quantification of (D) total $Ca^{2+}$ peaks per ovulation and (E) peak frequency (peaks/min). In (D, E), symbols denote transit failures: trapped oocytes (square) and pushback (star). (F) Representative 2D kymograms of spatiotemporal $Ca^{2+}$ during independent ovulations across indicated genotypes. Panel borders are color-coded by final transit outcome: successful exit (black), pushback (orange), or trapped (blue). The black arrow indicates an abnormal event where two oocytes entered the spermatheca and were both pushed back. Data are presented as mean ± SD. Statistical significance was determined using an ordinary one-way ANOVA followed by Dunnett's multiple comparisons test (*$p < 0.05$, **$p < 0.01$, ***$p < 0.001$, ****$p < 0.0001$, ns = not significant).

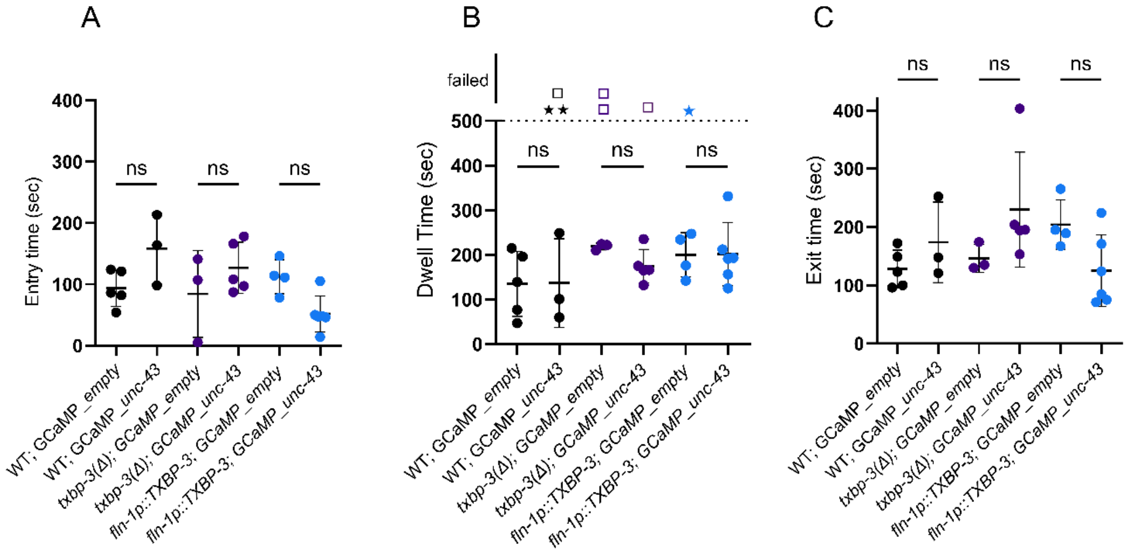


**Figure S3. Depletion of *unc-43* by RNAi does not significantly affect oocyte transit times in the GCaMP3 background.** (A-C) Quantification of oocyte (A) entry, (B) dwell, and (C) exit times in GCaMP-expressing WT, *txbp-3(Δ)*, and *fln-1p::*TXBP-3 animals treated with *empty*(RNAi) or *unc-43*(RNAi). In (B), symbols denote transit outcomes: successful (solid circles, <500 sec) and failed transits (>500 sec), including trapping (squares) and pushback (star). Data are presented as mean ± SD. Statistical significance was determined using a one-way ANOVA followed by Šídák's multiple comparisons test (ns = not significant).

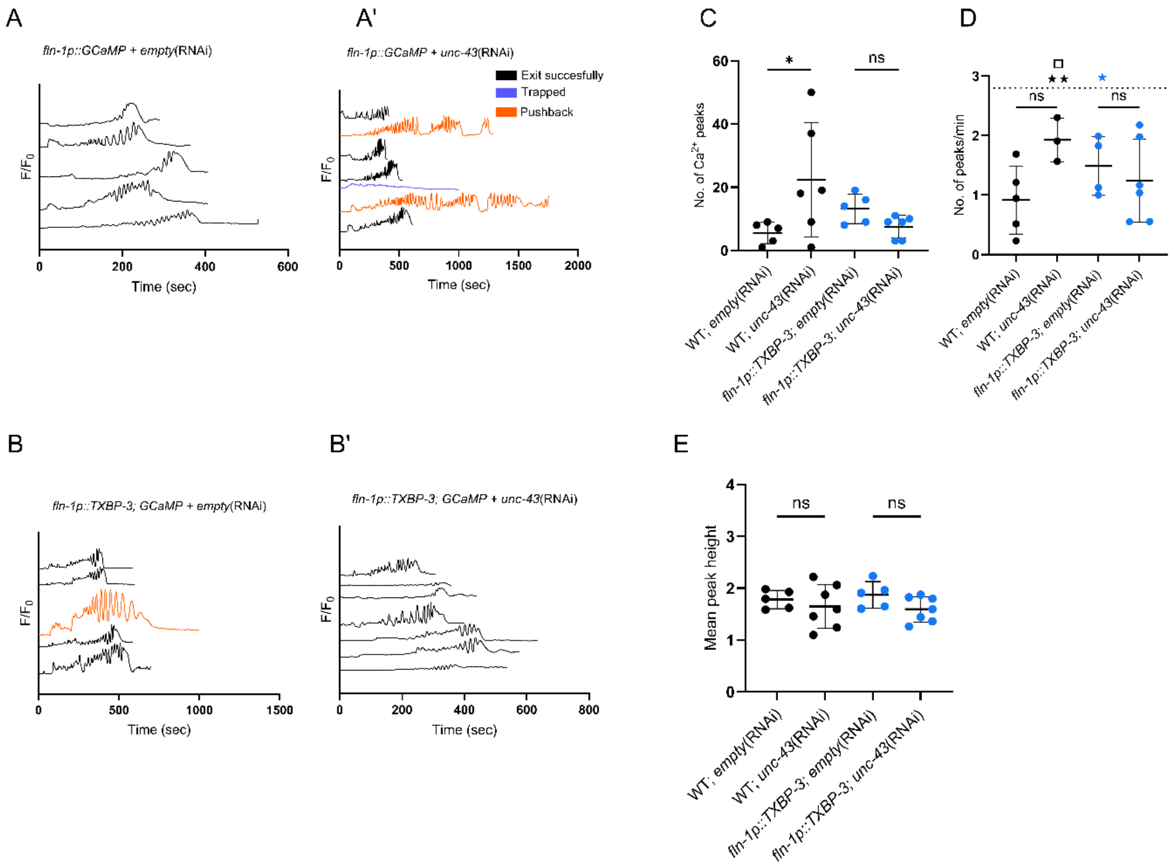


F. *fln-1p::GCaMP; empty*(RNAi)

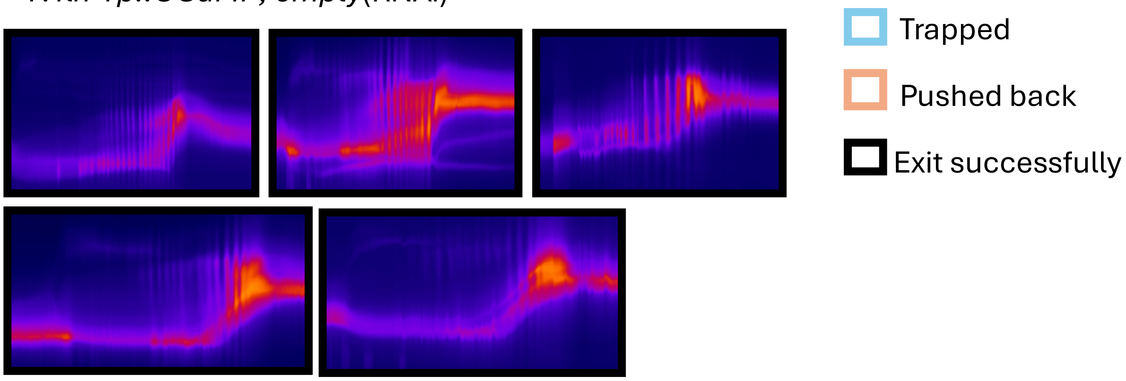

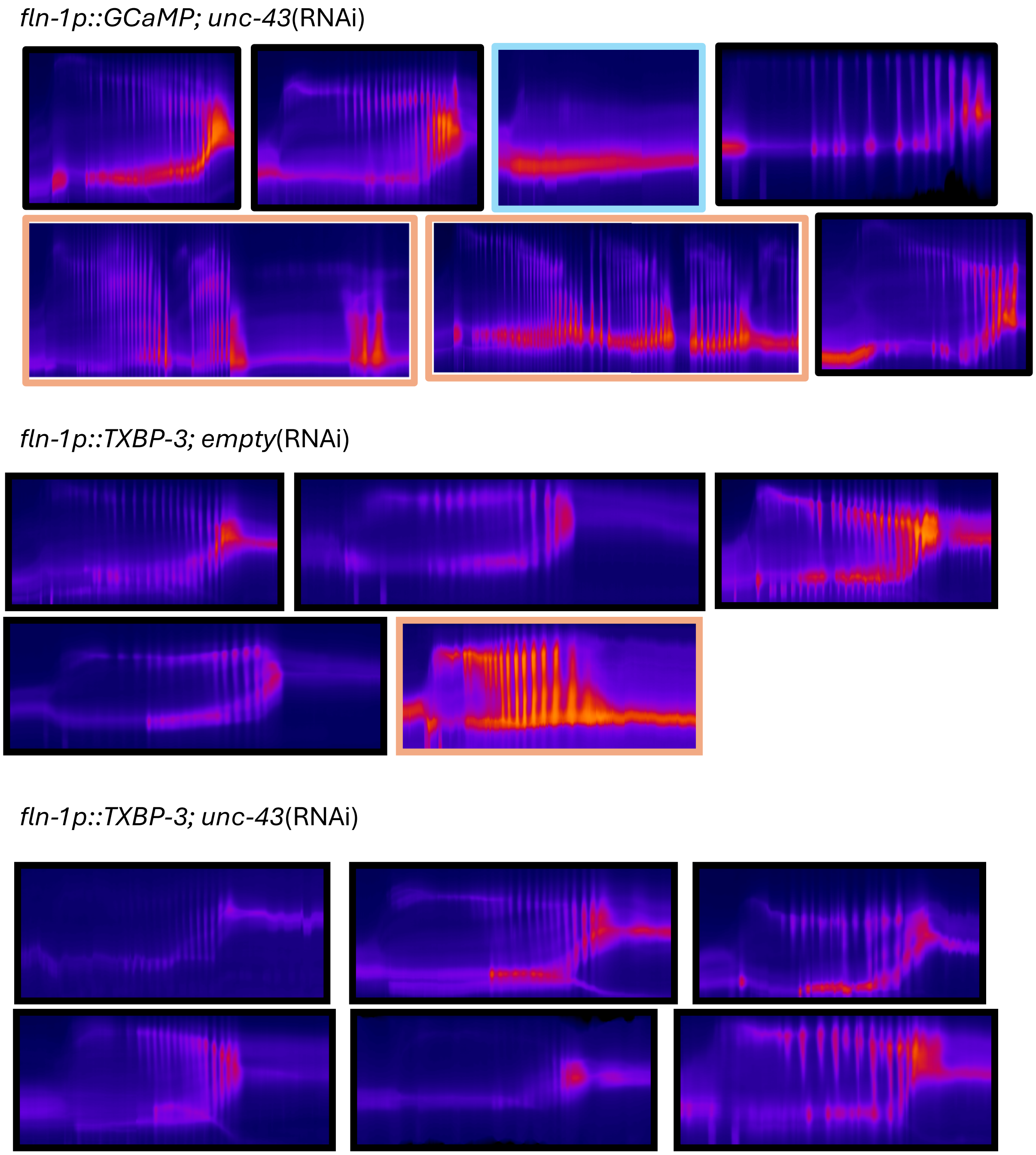


**Figure S4. Effect of *unc-43*(RNAi) on spermathecal $Ca^{2+}$ dynamics.** (A-B') Normalized 1D $Ca^{2+}$ traces for (A) WT; *empty*(RNAi), (A') WT; *unc-43*(RNAi), (B) *fln-1p::*TXBP-3; *empty*(RNAi), and (B') *fln-1p::*TXBP-3; *unc-43*(RNAi). Traces are color-coded by transit outcome: successful exit (black), trapped (blue), and pushback (orange). (C-E) Quantification of (C) total $Ca^{2+}$ peaks per transit, (D) peak frequency (peaks/min), and (E) mean peak height. In (D), symbols above the dotted line denote transit failures: trapped oocytes (square) and pushback into the gonad arm (star). (F) Representative 2D kymograms of individual transits across WT and *fln-1p::*TXBP-3 backgrounds treated with *empty*(RNAi) or *unc-43*(RNAi). Kymogram borders indicate mechanical outcome: successful exit (black), trapped (blue), or pushback (orange). Data are presented as mean ± SD. Statistical significance was determined using an ordinary one-way ANOVA followed by Šídák's multiple comparisons test (*$p < 0.05$, ns = not significant).
